# Disentangling the effects of Anxious, Autistic and Schizotypal Traits on Perceptual Inference

**DOI:** 10.1101/2024.10.14.618266

**Authors:** Caroline Bévalot, Maia Mallevays, Emile Bayon de Noyer, Lucie Berkovitch, Florent Meyniel

## Abstract

Perception combines sensory information with prior knowledge, but people differ in how they do so. These differences may be associated with psychological traits; however, previous studies often focused on one trait at a time and yielded contradictory results. We measured how anxious, autistic and schizotypal traits relate to perception in two general population samples (original: n=277; replication: n=279, pre-registered). We combined tasks in which priors were either explicitly communicated or implicitly learnt with computational modelling to distinguish the decision and learning stages of perceptual inference. The three traits formed distinct but correlated dimensions. At the decision stage, higher trait anxiety was associated with more optimal choices and greater reliance on sensory information, without any alteration in the reliance on priors, whether explicit or implicit. On the other hand, state anxiety was associated with a reduced effect of sensory information on response speed. When traits were analysed separately, autistic traits showed similar associations, but these associations disappeared once anxiety was taken into account in multidimensional analyses. No other credible or consistent effects emerged for autistic or schizotypal traits, nor for learning parameters. These results highlight the importance of a multidimensional approach, including anxiety measurements, to characterise inter-individual differences in perception.

## 1. INTRODUCTION

Our brain is constantly confronted with uncertainty when perceiving its environment. On the one hand, the sensory information it receives is noisy and ambiguous. On the other hand, the prior representations (i.e., prior beliefs about the environment) that are used to compensate for the limited sensory information are also uncertain and subject to change in a dynamic environment [1–3].

Bayesian inference provides an optimal solution for combining sensory information (known mathematically as “likelihood”) with priors, and for learning these priors from past observations [4]. Perceptual inference includes two distinct stages. At the perceptual decision stage, Bayes’ rule prescribes that in an optimal observer, the posterior probability of a given interpretation for a sensory input is the product of its prior probability (i.e., how probable that interpretation is before observing the input) and its sensory likelihood (i.e., the sensory evidence in its favour). These priors must be learned: at the learning stage, they are acquired and updated from past observations, at a rate that should depend on the volatility of the environment (i.e., how frequently its underlying statistics change). Humans follow Bayesian inference to some extent in perception [5–8], as shown notably in early multisensory integration studies [9,10]. However, the extent to which they depart from Bayesian computations (i.e., optimality) varies considerably across individuals (see [11] for a review).

Interindividual differences in perceptual inference have been associated with psychological traits, but the literature reports contradictory findings. Many Bayesian accounts of psychotic manifestations (from schizotypal traits in the general population to psychotic symptoms in clinical populations) have been proposed (reviews: [12–20]). However, the evidence remains mixed: psychosis has been associated with a precision of priors that is either abnormally high [21–27] or abnormally low [12,28–32]. Similarly, some researchers have proposed Bayesian accounts of autism, with mixed results (reviews: [33–38]). Indeed, autistic traits have been proposed to result from an abnormally weak influence of priors [37], from an excessive influence of sensory likelihood [34], from an excessive weight of prediction errors [38], and from an overestimation of the volatility of the environment [36]. Finally, anxious traits have been described through the lens of Bayesian inference (reviews: [39–45]). Again, the Bayesian accounts vary: anxiety has been related to an altered processing of uncertainty [40,42,45], a misinterpretation of stochasticity as volatility [44] and an increased reliance on priors [46].

These inconsistencies regarding the relationship between psychological traits and perceptual inference may stem from methodological differences across studies, including experimental design, modelling, and analysis. First, previous studies differ in whether their designs investigate the prior-likelihood combination, the learning of a prior, or both. Not distinguishing these two aspects of perceptual inference makes conclusions ambiguous [26,47–56]. For instance, an apparent overweighting of priors in decision could actually result from a prior value more extreme than it should be due to abnormal learning, with no alteration of its weighting during the prior-likelihood combination at the decision stage.

Second, modelling choices can limit the distinction between several alterations. Specifically, the weights of the prior and the sensory likelihood onto perceptual decisions can be captured as a single relative weight (or balance) between the two [23,52,53,57–59], or two separate weights [48,49,54], and the first option (a relative weight) cannot distinguish the weights of the prior and the likelihood.

Third, beyond experimental and modelling differences, psychological traits are analysed differently across studies. Many studies adopt a categorical approach [23,48,54,60,61], dividing participants based on a clinical threshold. However, psychological traits lie on a continuum rather than forming clusters [62,63]. In addition, most previous studies focused on a single psychological trait, e.g. psychotic [26,51–53,56], autistic [47,50,55,59], anxious [57,64]. Yet these traits are often strongly correlated: anxiety disorders affect 45% of people with schizophrenia and 39.6%-55.3% of those with autism [65,66], against only 16% in controls [67]; schizophrenia is 3.6 times more prevalent in autism than in controls [68]. Overlooking these correlations is particularly problematic if traits are associated with opposing differences in perceptual inference. A unidimensional approach could miss trait-specific effects or spuriously attribute an effect to the wrong dimension.

In the current study, we specifically designed cognitive tasks and modelling approaches to overcome these limitations. First, we distinguished perceptual decision-making from learning within the same subjects by using two tasks which differed in the way priors are provided to participants: either explicitly or implicitly. Priors are considered explicit when they are directly communicated to participants, and implicit when participants have to learn them from observations. Perceptual decisions must combine likelihoods and priors in both cases, but learning is required only with implicit priors. Second, we modelled the weights of the prior and the likelihood separately. We used a Bayesian model capturing departures from optimality separately at each stage (Figure 1): at the decision stage, participants may over- or underweight the prior and the sensory likelihood independently or show response biases; at the learning stage, they may misestimate the volatility of the environment, or make an exacerbated, dampened or biased use of the sensory likelihood when updating their priors. Finally, we used a multidimensional approach to analyse psychological traits’ influence on perceptual inference, which characterises a continuum of anxious, autistic, and schizotypal traits simultaneously. To reinforce the robustness of these findings, we collected two independent datasets and checked the reproducibility of the results.

**Figure 1.**
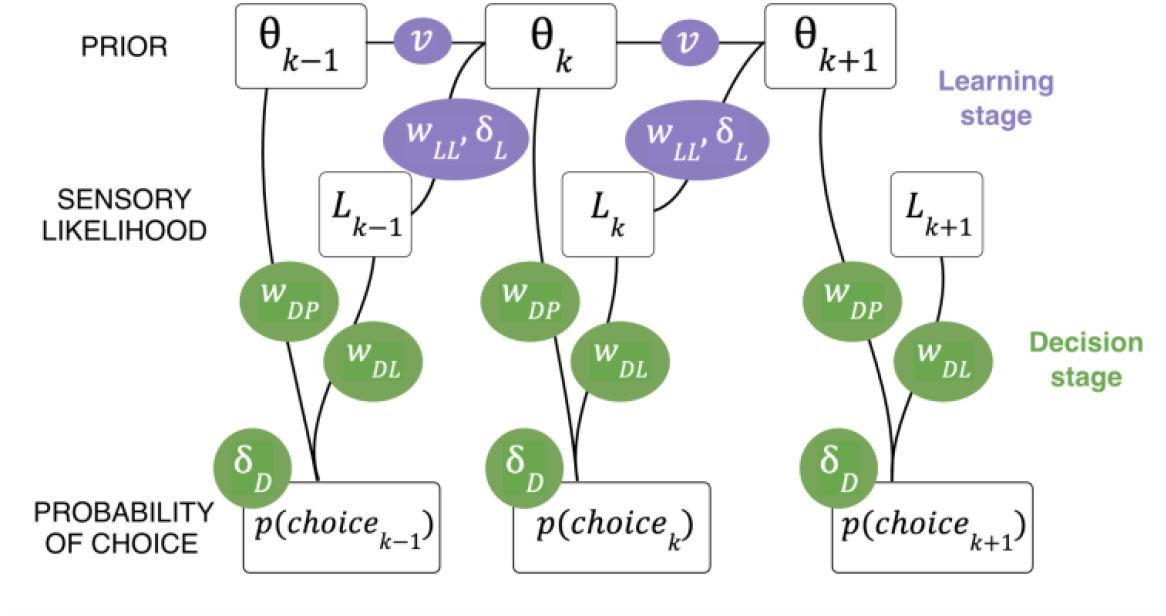
Schematic of the information flow in the Bayesian model of learning and perceptual decision making. The variables in the ovals correspond to the free parameters of learning (purple) and decision making (green) stages. Choices follow probabilistically a combination of prior probability and sensory likelihood (with weighting factors *W_DP_* and *W_DP_*, respectively, and a bias *δ_D_*). The learned prior value θ is updated based on the sensory likelihood (with a bias *δ_L_*,and weighting factor *W_LL_*) by assuming a volatility level (i.e. change point probability) *v*.

## 2. METHODS

See Supplementary Materials for full methods.

### 2.1 Participants

The study was approved by the local Ethics Committee (CER Paris Saclay, n°222) and participants gave their informed written consent before participating. Participants were recruited online via Prolific.co and compensated for their time. Data were collected in two independent samples: September 2021 (Original Sample: S1) and March 2023 (Replication Sample: S2). S1 comprised 277 participants (136 female, 141 male; age range: 19-53, mean=31, SD=8) and S2 comprised 279 participants (132 female, 143 male, 4 unreported; age range: 19-53, mean=33, SD=9). Completion of both tasks and all psychometric questionnaires was required for participation to be validated. Participants were excluded if they responded on fewer than 25% of trials, resulting in final samples of N=274 (S1) and N=279 (S2). As sex was included as a covariate in the regression models, the 4 participants with unreported sex in S2 were not included in the regression analyses (N=275). Results are presented separately for the two independent samples.

### 2.2 Psychological trait questionnaires

Participants completed questionnaires assessing three psychological traits: anxiety (State Trait Anxiety Inventory, STAI; [69,70]), autistic traits (Autistic Quotient, AQ; [71,72], and schizotypal traits (Schizotypal Personality Questionnaire, SPQ; [73–75]). The STAI distinguishes between state anxiety (STAI-A), reflecting immediate emotional reactions, and trait anxiety (STAI-B), a stable dispositional characteristic; we included both to disentangle enduring psychological traits of anxiety from transient anxiety during the experimental assessment. The SPQ captures three dimensions of schizotypal traits: a cognitive-perceptual dimension characterised by unusual perceptual experiences, magical ideation, and ideas of reference (which can be related to positive symptoms); an interpersonal dimension characterised by social withdrawal, reduced emotional expression, and social anxiety (which resembles negative symptoms); and a disorganisation dimension characterised by odd speech and odd behaviour [75]. Given that alterations in Bayesian perceptual inference are most consistently associated with positive psychosis symptoms [25,27,76,77], we focused our main analyses on the cognitive-perceptual dimension, with interpersonal and disorganisation results presented in Supplementary Materials.

### 2.4 Tasks: using explicit and implicit priors to study two stages of perceptual inference

We used two separate tasks to study interindividual differences and distinguish two stages of perceptual inference: decision and learning (Figure 2). These tasks and the model have previously been used by Bévalot and Meyniel [78] to characterise perceptual inference. Subjects had to categorise as house or face each noisy morphed image presented in a sequence. For each image, the likelihood of the category was estimated empirically as the proportion of human observers in an independent sample who categorised this image as a house rather than a face (see [78]). On a given trial, the image category was sampled with probability 0.1, 0.3, 0.5, 0.7 or 0.9 to be a house, which we call generative prior hereafter.

**Figure 2.**
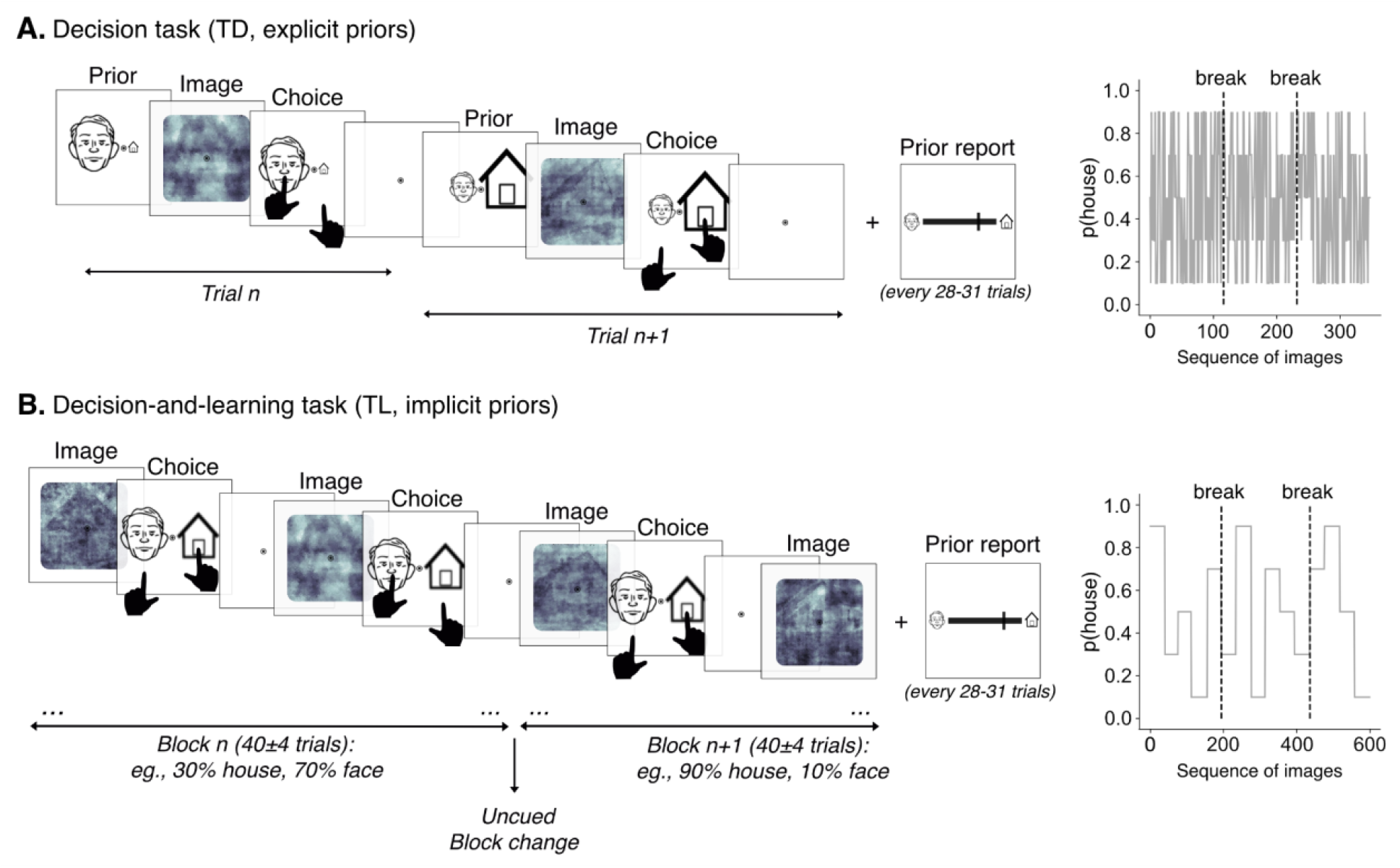
Learning and perceptual decision-making tasks. Subjects categorised each noisy image as face or house. The sensory likelihood conveyed by each image was measured in an independent group of subjects. Priors correspond to the prior probability of the latent category (face or house) on each trial. (A) In the decision task, priors changed from trial-to-trial and were communicated explicitly as the relative size of pictograms presented before each image (and reminded to subjects at the response stage). (B) In the learning-and-decision task, priors were not communicated but they remained constant within uncued blocks of 40±4 trials, making it possible to learn their value from the sequence of images. Subjects were occasionally asked to report the prior value. Right panel: Sequence of generative prior values used in (A) the decision task, and in (B) the decision- and-learning task for an example participant. In both tasks, the sequence is split in three sessions separated by short breaks.

In the “decision task” (including decision only, TD), priors about the category were provided explicitly before each image in the form of pictograms. Subjects were instructed that the relative size of the house and face pictograms denoted the prior value (see Figure S1 for instruction slides). In the “decision-and-learning task” (including decision and learning, TL), such explicit cues were not provided; instead, subjects were instructed that the implicit prior was constant in uncued blocks of trials, making it possible for them to learn the prior value (Figure 1B) (see Figure S2 for instruction slides). By contrast, in the “decision task”, explicit priors changed from trial to trial, preventing learning. See Supplementary methods for further details. We report results separately for both tasks (TD and TL) and for both samples (S1 and S2). For example, TLS1 corresponds to the decision-and-learning task in the original sample.

### 2.5. Using Bayesian inference to quantify parameters of perceptual decision and learning

#### 2.5.1. Optimal Bayesian observer: logistic regression weights of sensory likelihood and priors

We characterised perceptual inference using a model derived from Bayesian inference to analyse participants’ choices, distinguishing two stages: the decision stage (for both tasks) and the learning stage (for the decision-and-learning task only). At the decision stage, Bayes’ rule prescribes that the posterior probability of the latent category, which determines the agent’s choice, is the product of the prior and the sensory likelihood (Figure 1). In log-odds form this combination is additive (log-posterior-odds = log-prior-odds + log-likelihood-ratio; see Supplementary Methods Eq. 1-3), which motivates using logistic regression to quantify each term’s separate contribution to choice. Note that in the decision task, as priors were provided explicitly before each image, no learning is required, and we used the generative prior value in the regression analysis. In contrast, in the decision-and-learning task, the prior is implicit as it must be inferred over time from previous observations in uncued blocks of trials. To capture how these implicit priors can be learned ideally from previous observations, we applied a BAYES-OPTIMAL model which inverts exactly the generative process of the sequence of images (Supplementary Methods Eq. 4-6).

#### 2.5.2. Parameterised Bayesian observer: fitted weights of decision and learning parameters

Human participants may depart from the BAYES-OPTIMAL model in several ways (Figure 1). At the perceptual decision stage, they may over or underweight the prior and/or sensory likelihood or show response biases. At the learning stage, participants may misestimate how frequently the generative priors change [79,80], or make an exacerbated, dampened or biased use of the sensory likelihood when updating their beliefs. To quantify these biases, we used the BAYES-FIT-ALL model, previously designed and validated [81]. This model is a parameterised version of the Bayesian observer, which we fit to each participant’s choices by maximum likelihood, allowing us to extract participant-specific parameters of perceptual decision and learning (Supplementary Methods Eq. 7). The model has six free parameters: three for decision (weight of prior on choices, weight of likelihood on choices, decision bias) and three for learning (volatility, weight of likelihood in learning of priors, learning bias) (Figure 1).

#### 2.5.3. Reaction time

We also analysed the effects of parameters of perceptual decision and learning on response speed (the inverse of reaction time) in linear regressions (one regression per factor). Choices and reaction times reflect complementary behavioural readouts of perceptual inference. Choices depend on the value of the prior and the likelihood, i.e., which category they point toward. Response speed, by contrast, does not depend on which category is favoured, but how strongly the prior and the likelihood favour one category (no matter whether it is a face or house). Therefore, to analyse response speed, we used the strength of likelihood (Supplementary Methods Eq. 9), and the strength of the prior (Eq. 10). We also tested whether confidence about the prior value (i.e., its posterior log-precision as learned by the BAYES-OPTIMAL model; Eq. 11) could influence response speed. Confidence about the prior value should be used in the learning stage, to determine how much the current prior is trusted when computing the next one, which may be reflected in response speed. For this analysis only, trials with RTs < 100 ms or >1950 ms were excluded (TDS1: 8%; TLS1: 7.6%; TDS2: 3.6%; TLS2: 2.4% of trials).

### 2.5.4. Multiple linear regression models to quantify associations with traits

In the previous sections, we have derived several parameters that characterize perceptual inference: the logistic regression weights of prior and likelihood on choices, the fitted parameters of decision and learning in the BAYES-FIT-ALL model, and the linear coefficients of sensory likelihood strength, prior strength and confidence about the prior onto response speed. We tested whether these parameters of perceptual inference could be associated with psychological traits and sociodemographic data, using a separate multiple linear regression for each perceptual inference parameter (Supplementary Methods Eq. 8).

## 3. RESULTS

### 3.1. Participants use both priors and sensory likelihood in their choices

At the group level, subject responses showed clear effects of the generative priors (reflected by the pictograms in the decision-task TD and the true prior value in the decision-and-learning task TL) and the sensory likelihood in both tasks (Figure 3, Table 1). Similarly, there was a significant effect of the optimal Bayesian prior (reflecting learning of implicit priors) in the TL in both samples (Table 2). This effect was stronger than the one of generative priors, which is consistent with the fact that generative priors are not provided to subjects in this task, they must be learned (S1: mean difference=0.12, std=0.14, t=14.14, p<10^−33^; S2: mean difference=0.13, std=0.11, t=18.84, p<10^−52^).

**Table 1.** Logistic regression weights of priors and sensory likelihood on choices (with generative or optimal priors) in the decision-only task (TD) and the decision-and-learning task (TL), in samples S1 and S2. We assessed whether these weights differed reliably from zero using Wilcoxon signed-rank tests for non-normally distributed variables. W = Wilcoxon signed-rank test; p = p-values (tests against a null of zero weight); d [95% CI] = Cohen’s d with 95% bootstrapped confidence intervals.

| Perceptual parameter | TaskSample | Mean weight ± SEM | *t*/*W* (p) | *d* [95% CI] |
|---|---|---|---|---|
| Prior (generative) | TDS1 | 0.82±0.06 | *W*=1166 (*p*<10<sup>-41</sup>) | 0.82[0.74, 0.90] |
|  | TDS2 | 0.85±0.05 | *W*=794 (*p*<10<sup>-44</sup>) | 1.00[0.90, 1.15] |
|  | TLS1 | 0.36±0.02 | *W*=1209 (*p*<10<sup>-41</sup>) | 1.37[1.16, 1.61] |
|  | TLS2 | 0.38±0.02 | *W*=974 (*p*<10<sup>-43</sup>) | 1.53[1.34, 1.76] |
| Likelihood<br>(with generative prior) | TDS1 | 0.54±0.03 | *W*=1318 (*p*<10<sup>-40</sup>) | 1.31[1.16, 1.48] |
|  | TDS2 | 0.68±0.02 | *W*=441 (*p*<10<sup>-45</sup>) | 1.93[1.67, 2.26] |
|  | TLS1 | 0.65±0.03 | *W*=1023 (*p*<10<sup>-42</sup>) | 1.40[1.22, 1.62] |
|  | TLS2 | 0.82±0.02 | *W*=517 (*p*<10<sup>-45</sup>) | 2.14[1.81, 2.60] |
| Prior (optimal) | TLS1 | 0.48±0.02 | *W*=1224 (*p*<10<sup>-40</sup>) | 1.30[1.10, 1.53] |
|  | TLS2 | 0.51±0.02 | *W*=831 (*p*<10<sup>-42</sup>) | 1.53[1.34, 1.76] |
| Likelihood<br>(with optimal prior) | TLS1 | 0.70±0.03 | *W*=965 (*p*<10<sup>-42</sup>) | 1.45[1.26, 1.67] |
|  | TLS2 | 0.87±0.02 | *W*=528 (*p*<10<sup>-45</sup>) | 2.19[1.84, 2.68] |

**Table 2.**
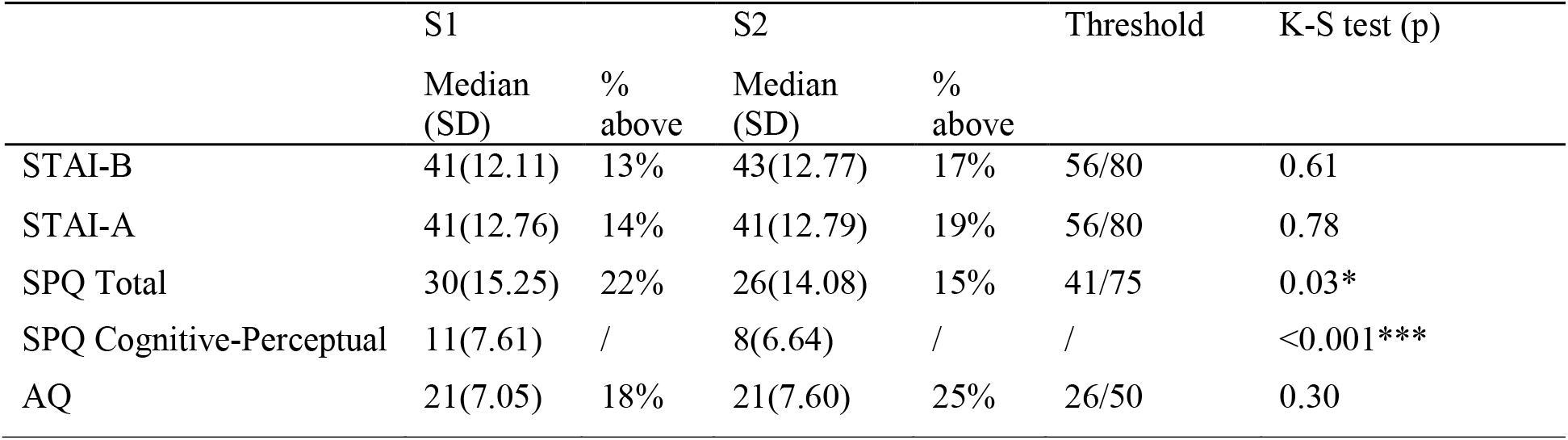
Trait medians and percentages above threshold (for considering high symptoms in healthy subjects) in original (S1) and replication (S2) samples. Kolmogorov-Smirnov (K-S) tests were used to check whether distributions were significantly different between samples S1 and S2 (***p<0.001, **p<0.01, *p<0.05). STAI-B = Trait Anxiety Inventory; STAI-A = State Anxiety Inventory; SPQ = Schizotypal Personality Questionnaire; AQ = Autistic Quotient.

**Figure 3.**
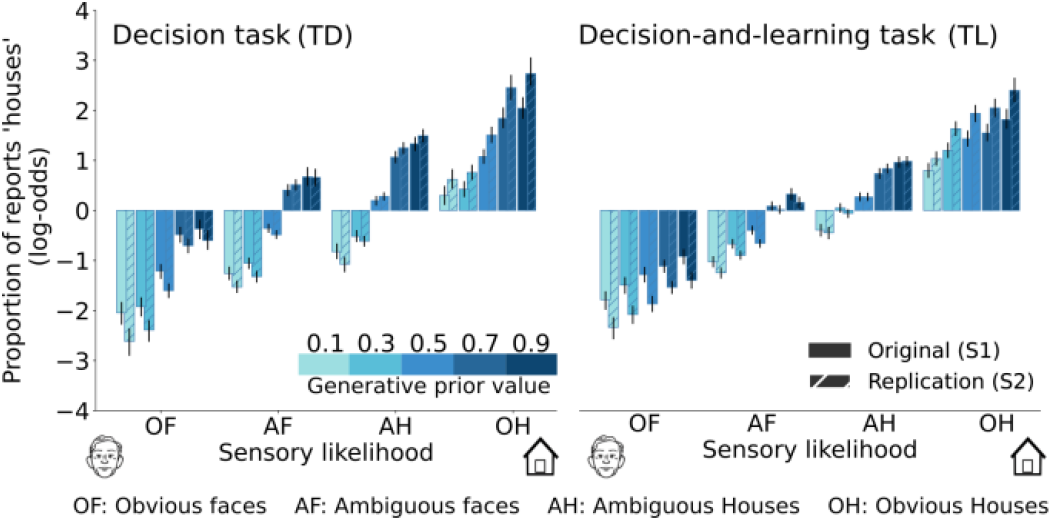
Priors and sensory likelihood influence perception whether priors were explicit or implicit. Average proportion of answers (plotted in log-odds) sorted by bins of sensory likelihood and generative prior values.

### 3.2. Anxious, autistic, and schizotypal traits correspond to three dimensions which are strongly correlated

We checked the quality of the self-reported questionnaire (STAI, SPQ, AQ) scores using Cronbach’s alpha, which assesses internal consistency of questionnaires. High alpha values were observed across both samples, consistent with those reported in original validation studies suggesting that the scores were reliable (Table S1; Figure S4).

Before adopting a multidimensional approach, we tested in a data-driven manner whether the three traits actually correspond to distinct psychological dimensions using a factor analysis. Such analysis operates on items ignoring the questionnaires they belong to; yet a three-factor analysis resulted in groups of items corresponding almost perfectly to the questionnaires. When four factors were used in the analysis (one more than the number of questionnaires), one questionnaire was split into two factors, namely the AQ in S1, and SPQ in S2 (Figure S5), so that the fourth factor did not isolate a fourth dimension compiling items from different questionnaires. Therefore, the STAI, AQ and SPQ questionnaires correspond to three separable dimensions, and thus, traits.

Although psychological traits were separable, we found them to be correlated (Figure 4A), similarly to previous studies, stressing the need for a multidimensional analysis, in which all traits are included in the same model to isolate the unique contribution of each. In each sample, subjects covered a wide range of psychological trait scores, allowing to analyse interindividual differences (Figure 4B). Distributions were largely comparable between samples, with the non-intended exception of schizotypal traits (Table 2). The original sample had significantly higher cognitive-perceptual and disorganization traits than the replication sample (Figure S6).

**Figure 4.**
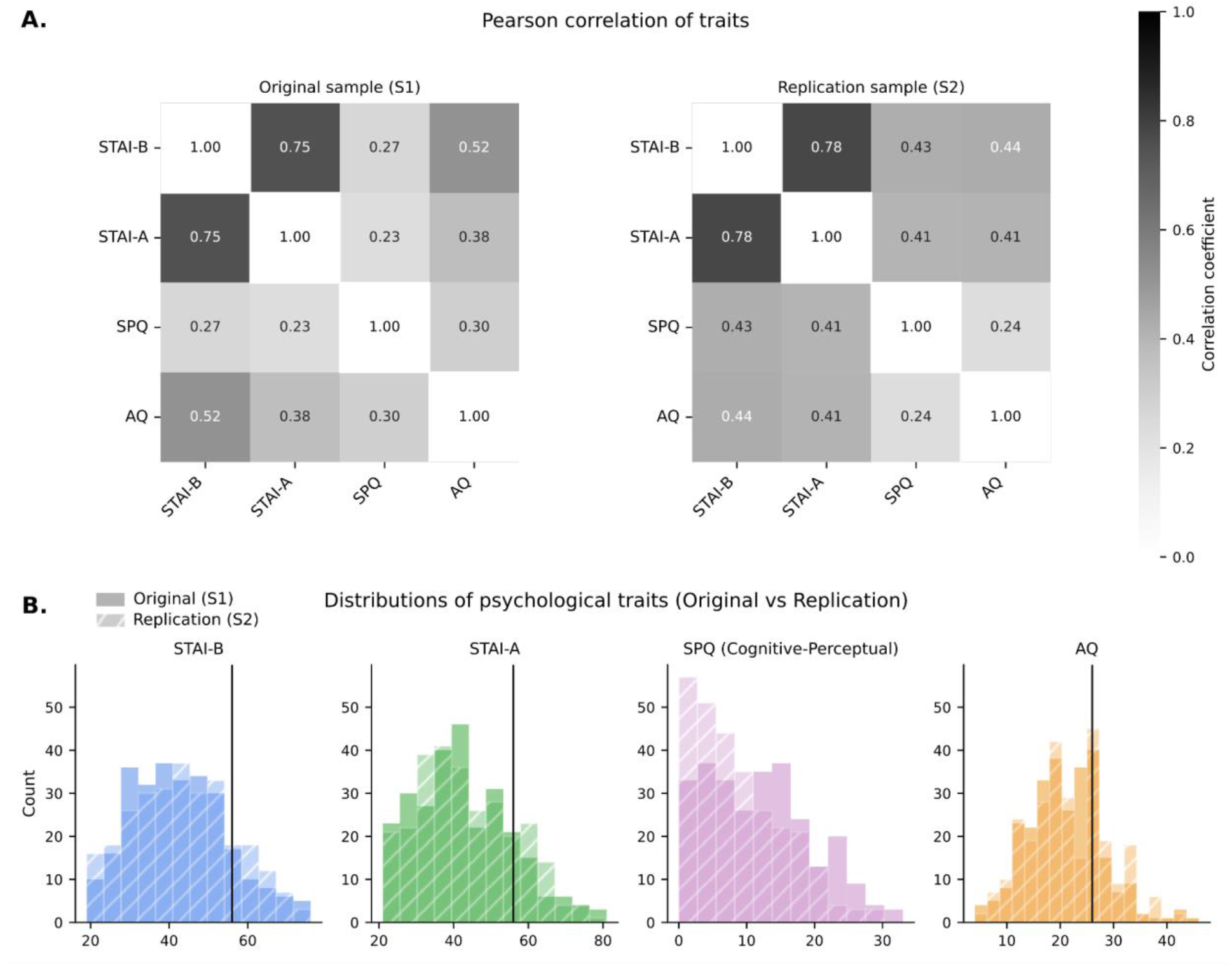
Psychological traits cover a broad range in both samples and are correlated to one another. A.Pearson correlation coefficient of the different traits. B.Distributions of each psychological trait in the original sample (solid) and the replication sample (hatched). Black vertical lines mark the clinical threshold for elevated symptoms in healthy subjects: 56 for STAI-A/B and 26 for AQ. For SPQ, there is no threshold because this threshold is defined for the total score (not shown) whereas we focused on the cognitive-perceptual dimension of the scale.

### 3.3. Anxiety is associated with more optimal choices

We tested whether psychological traits could be associated with differences in perceptual inference. We started by comparing participants’ choices to the posterior probability of the house/face category (in short “the posterior”). According to Bayes’ rule, this posterior probability corresponds to the statistically optimal choice given the sensory evidence and prior knowledge, the latter being the generative priors in TD or the priors inferred by the BAYES-OPTIMAL model in TL. In a logistic regression, a subject whose choices are more optimal (i.e. closer to the BAYES-OPTIMAL model) has a larger weight on the BAYES-OPTIMAL posterior (this weight should increase up to 1 for a fully optimal subject). We found that higher trait anxiety (STAI-B) was associated with more optimal choices, with positive coefficients in all samples which were significant in the TL in both samples (TLS1: p=0.010; TLS2: p=0.039), marginal in TDS1 (p=0.064), and non-significant in TDS2 (p=0.21)

(Figure 5A, Table S2). We observed the opposite pattern for state anxiety (STAI-A), but it did not reach significance (all ps>0.05, see Figure 5A, Supplementary Table S2). Results regarding the cognitive-perceptual dimension of schizotypal traits were inconsistent across samples (see Supplementary Figure S7 for the interpersonal and disorganisation dimensions) whereas autistic traits were not associated with any difference (Figure 5A, Table S2).

**Figure 5.**
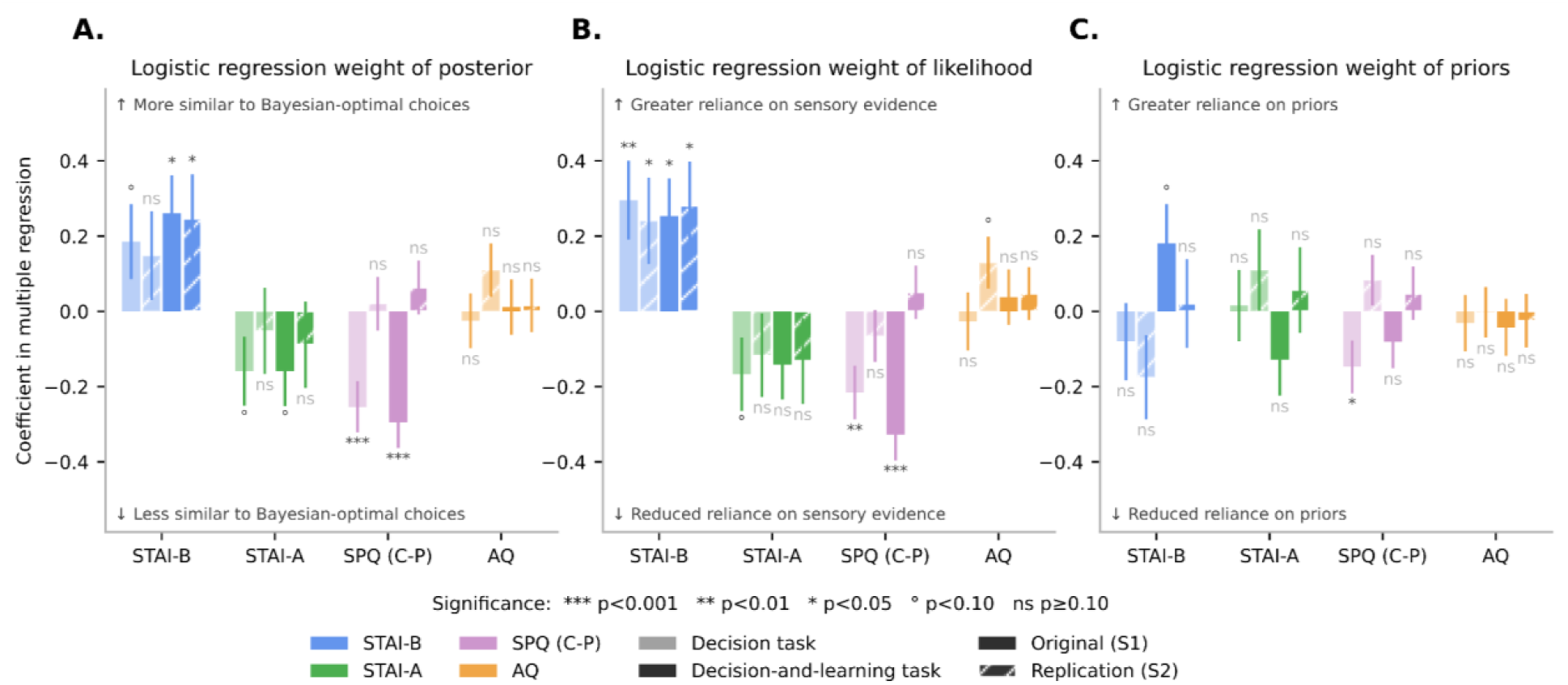
Anxious traits are associated with more optimal performance and greater weight on sensory likelihood. Coefficients (± SE) were derived from multiple standard linear regressions of logistic regression weights of perceptual parameters onto psychological traits and sociodemographic factors, estimated separately for each task and each sample. STAI-B = Trait Anxiety Inventory; STAI-A = State Anxiety Inventory; SPQ (C-P) = Schizotypal Personality Questionnaire (Cognitive-Perceptual dimension); AQ = Autistic Quotient. **A.Posterior weight**. In a Bayesian-optimal observer, choices are determined by the posterior probability. Therefore, positive coefficients indicate that people with higher trait levels make more Bayesian-optimal choices. **B.Likelihood weight**. Coefficients reflect the association between psychological traits and the influence of sensory evidence (likelihood) on choices. Positive coefficients indicate that people with higher trait levels rely more on sensory evidence in their perceptual decisions. **C.Prior weight**. Coefficients reflect the association between psychological traits and the influence of prior beliefs on choices. Positive coefficients indicate that people with higher trait levels rely more on priors in their perceptual decision. Note that for each participant, the weights of the prior and likelihood onto choices are estimated in the same logistic regression. In all three panels, posterior and prior probabilities are derived from the BAYES-OPTIMAL model.

### 3.4. Trait anxiety impacts the weight of sensory likelihood (but not priors) in perception

As priors and sensory likelihood jointly contribute to the posterior, we then studied how psychological traits would influence these two parameters separately (Figure 5B and C). Results are presented with Bayes-optimal priors, but we found similar results with BAYES-FIT-ALL priors (Figure S7).

Higher trait anxiety (STAI-B) was associated with greater reliance on sensory likelihood across both samples no matter the format of prior – explicit or implicit (Figure 5, Table S2). The impact of trait anxiety was specific to likelihood, as we found no significant associations between trait anxiety and the weight of explicit priors (TDS1, TDS2) or implicit priors (TLS1, TLS2). The bootstrapped difference between the regression coefficient of anxiety on likelihood weight versus prior weight confirmed this specificity when priors were explicit (TDS1: Δ=0.39, 95% CI [0.04, 0.74], p=0.034; TDS2: Δ=0.42, 95% CI [0.09, 0.74], p=0.018). The same pattern was observed with implicit priors in the TL, although it did not reach statistical significance (TLS1: Δ=0.08, 95% CI [-0.16, 0.31], p=0.53; TLS2: Δ=0.24, 95% CI [-0.079, 0.60], p=0.12). Finally, we did not find a significant effect of state anxiety (STAI-A) on the weight of sensory likelihood or priors on choices (Table S2).

Autistic traits were not associated with differences in the weight of the likelihood or of the prior on choices (Figure 5, Table S2). Note that a unidimensional analysis would have concluded that autistic traits are associated with greater weight of sensory likelihood on choices in the replication sample (TDS2: 0.19±0.06, p=0.0011; TLS2: 0.14±0.06, p=0.019), an effect which vanishes in a multidimensional analysis (see Figure S8 for complete comparison of uni- and multidimensional analysis).

Regarding schizotypal traits, results were inconsistent across samples, as seen in the analysis of the posterior. Stronger schizotypal cognitive-perceptual traits were associated with lower reliance on sensory likelihood in the original sample (TDS1, TLS1), but not the replication sample (TDS2, TLS2) (Table S2, Figure 5). In exploratory analyses, we investigated the effect of all three SPQ dimensions, cognitive-perceptual, interpersonal, and disorganization, on perceptual parameters, but the latter two were also not associated with any significant or consistent differences (Figure S9). Lower cognitive-perceptual trait scores in the replication sample (Figure S7) may partly explain the lack of replication of effects in this sample (see also Figure S10).

To quantify the extent to which differences in traits contribute to differences in perceptual inference, we used a dominance analysis. Note that because traits are correlated, the coefficients of the multiple linear regressions presented above do not quantify the relative contribution of each trait in explaining inter-individual variance. Dominance analysis addresses this by averaging each predictor’s contribution to the explained variance across all possible sub-models. This analysis showed that trait anxiety contributed to differences in perceptual inference to an extent comparable to that of demographic factors, notably age (See Figure S11).

### 3.5. Anxiety accounts for inter-individual differences at the decision stage rather than at the learning stage

The TL combined with the BAYES-FIT-ALL model allowed us to distinguish between the weight of sensory likelihood during the decision stage (*W*_DP_) and its weight during prior learning (*W*_DL_). More anxious traits were associated with larger weights on sensory likelihood at the decision stage (TLS1=0.24±0.10, p=0.02; TLS2=0.28±0.12, p=0.018) (Figure 6), mirroring the pattern observed with the BAYES-OPTIMAL model in both tasks (Figure 5). Interestingly, we found no such association between anxiety and the weight of the sensory likelihood at the learning stage (TLS1=−0.001±0.10, p=0.99; TLS2=0.02±0.12, p=0.89), indicating that anxiety accounts for inter-individual differences in perceptual inference at the decision stage, and not in the learning of a prior. No other fitted parameter was consistently associated with anxious traits in either sample. Autistic traits were not associated with differences in any decision or learning parameters (all ps>0.13). Schizotypal traits showed the same inconsistent results across samples as in the previous logistic regression analysis regarding the weight of sensory likelihood at the decision stage, and no significant effect on any other parameter.

**Figure 6.**
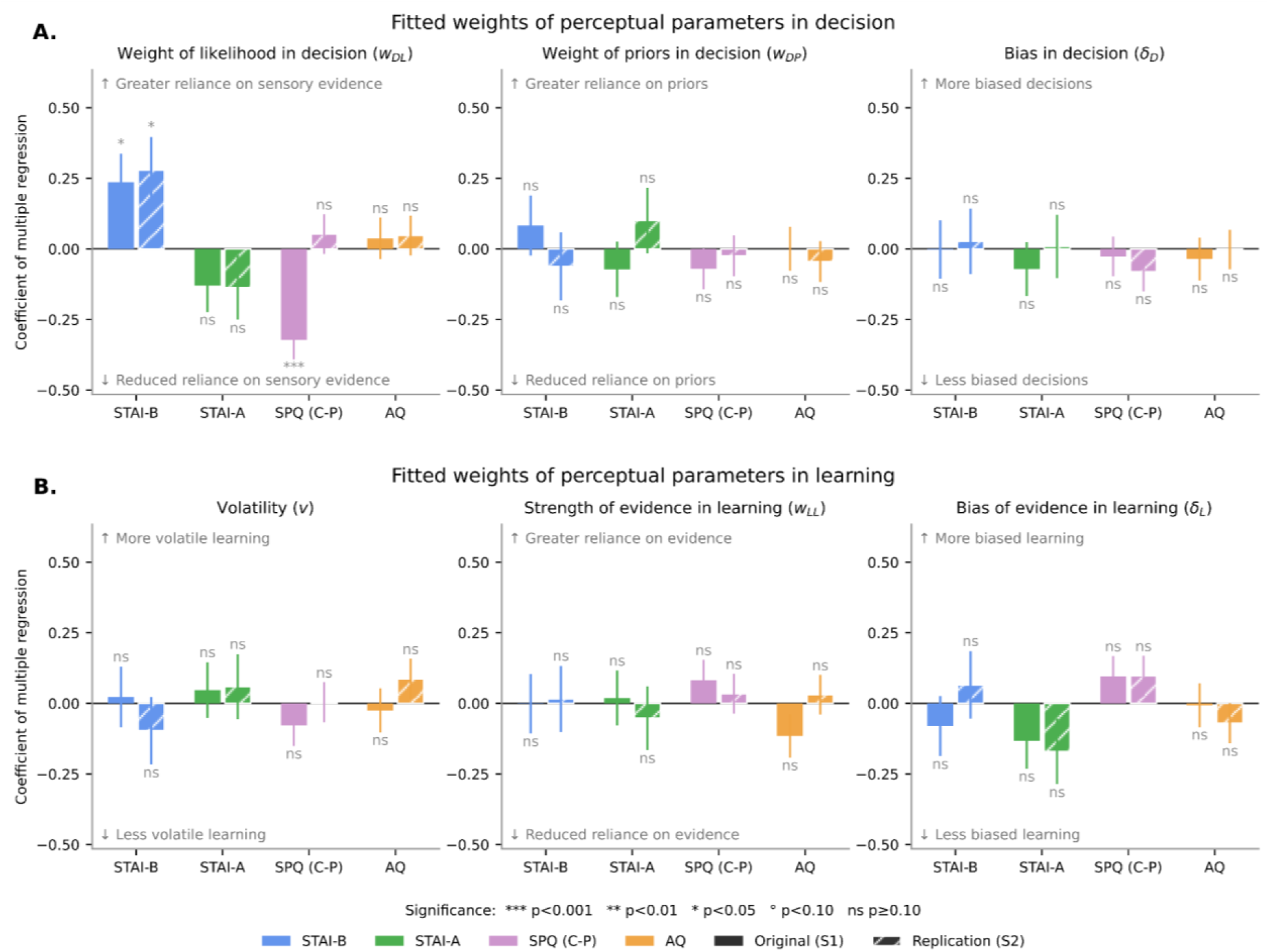
Associations between psychological traits and subject-specific parameters of perceptual inference fitted with the BAYES-FIT-ALL model. Anxiety traits are associated with an increased weight of likelihood on decisions; no consistent differences are observed for other traits or for learning parameters. Coefficients (± SE) were derived from multiple standard linear regressions of fitted model parameters onto psychological traits and sociodemographic factors, estimated separately for each task and each sample. Same plotting convention as in Fig 5. **(A)Decision parameters**. Weight of sensory likelihood in decision; weight of priors in decision; bias in decision. **(B)Learning parameters**. Volatility; strength of evidence in learning; bias of evidence in learning.

### 3.6. Reaction times confirm an association between trait anxiety and perceptual inference

To further validate our findings from the choice analysis, we examined whether the same underlying computations also account for interindividual differences in response speed (the inverse of reaction time). Response speed, unlike choice, should not depend on which category is favoured by the likelihood and the prior, but how strongly they support a category (whatever it is). Therefore, we used the strength of the likelihood (which quantifies how strongly the likelihood supports either category), and the strength of the prior (which quantifies how strongly either category is expected on the upcoming trial) as predictors. We used as an additional regressor the confidence about the prior value (i.e., its posterior log-precision as learned by the BAYES-OPTIMAL model). Confidence about the prior should influence the learning process (not which category is selected) and may therefore be reflected in the response speed. We found that subjects were faster when sensory likelihood was stronger, when the prior was stronger, and when confidence in the prior was larger (Table S3, Figure 7A).

**Figure 7.**
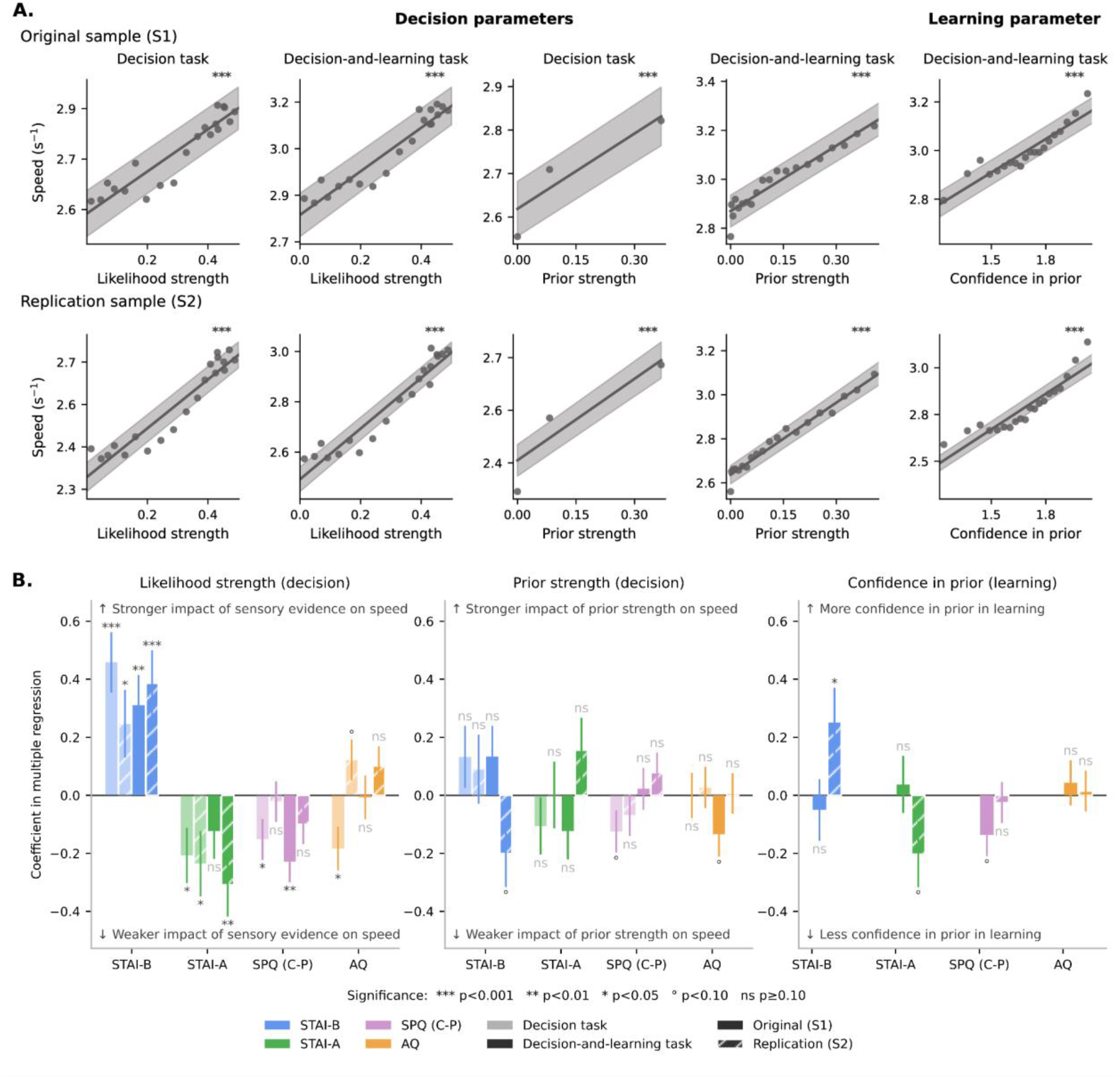
Effects on response speed corroborate those on choices. Correlations between response speed and sensory likelihood strength (eq. 9), prior strength (eq. 10), and confidence about the prior value (eq. 11), shown separately for the first (top row) and second samples (bottom row). Each scatter plot uses bins of values for visualization; statistics are computed on raw values across trials. Error shading corresponds to the SEM across subject-levels regression lines. ***p<0.001.

We then tested whether psychological traits could explain interindividual differences in how response speed was impacted by the likelihood strength, prior strength, and confidence. Anxious traits (STAI-B) were associated with an increased effect of likelihood strength, while we found no consistent association with prior strength or confidence (Figure 7B, Table 3). This association between anxious traits and the effect of likelihood strength was specific to likelihood strength relative to prior strength in TDS1 and TLS2 (bootstrapped difference between regression coefficient of anxiety on likelihood strength vs. prior strength: TDS1: Δ=0.32 [95% CI 0.03, 0.62], p=0.032; TLS2: Δ=0.58 [95% CI 0.22, 0.94], p=0.006), and the difference was in the same direction in TLS1 and TDS2 but did not reach statistical significance (TLS1: Δ=0.20 [95% CI −0.12, 0.5-], p=0.21; TDS2: Δ=0.17 [95% CI −0.15,0.51], p=0.32). In contrast to trait anxiety, state anxiety (STAI-A) was associated with a decreased effect of likelihood strength on response speed, which was significant in all samples except TLS1 (TDS1: p=0.032; TDS2: p=0.04; TLS1: p=0.20; TLS2: p=0.006, Figure 7B, Table 3). Trait anxiety (STAI-B) was also associated with an increased effect of confidence in prior on response speed in TLS2 (0.25±0.12, p=0.032), but this was not replicated in TLS1, where the coefficient was close to zero and in the opposite direction (TLS1: −0.05±0.12, p=0.64) (Figure 7, Table 3). Autistic traits were not associated with any significant effects (Table 3). For schizotypal traits, results were associated with reduced impact of strength of evidence in the first sample (TDS1), but not significantly so in the second (TDS2). Schizotypal traits were not associated consistently and significantly with any differences in the effects of prior strength or confidence (Table 3).

**Table 3.**
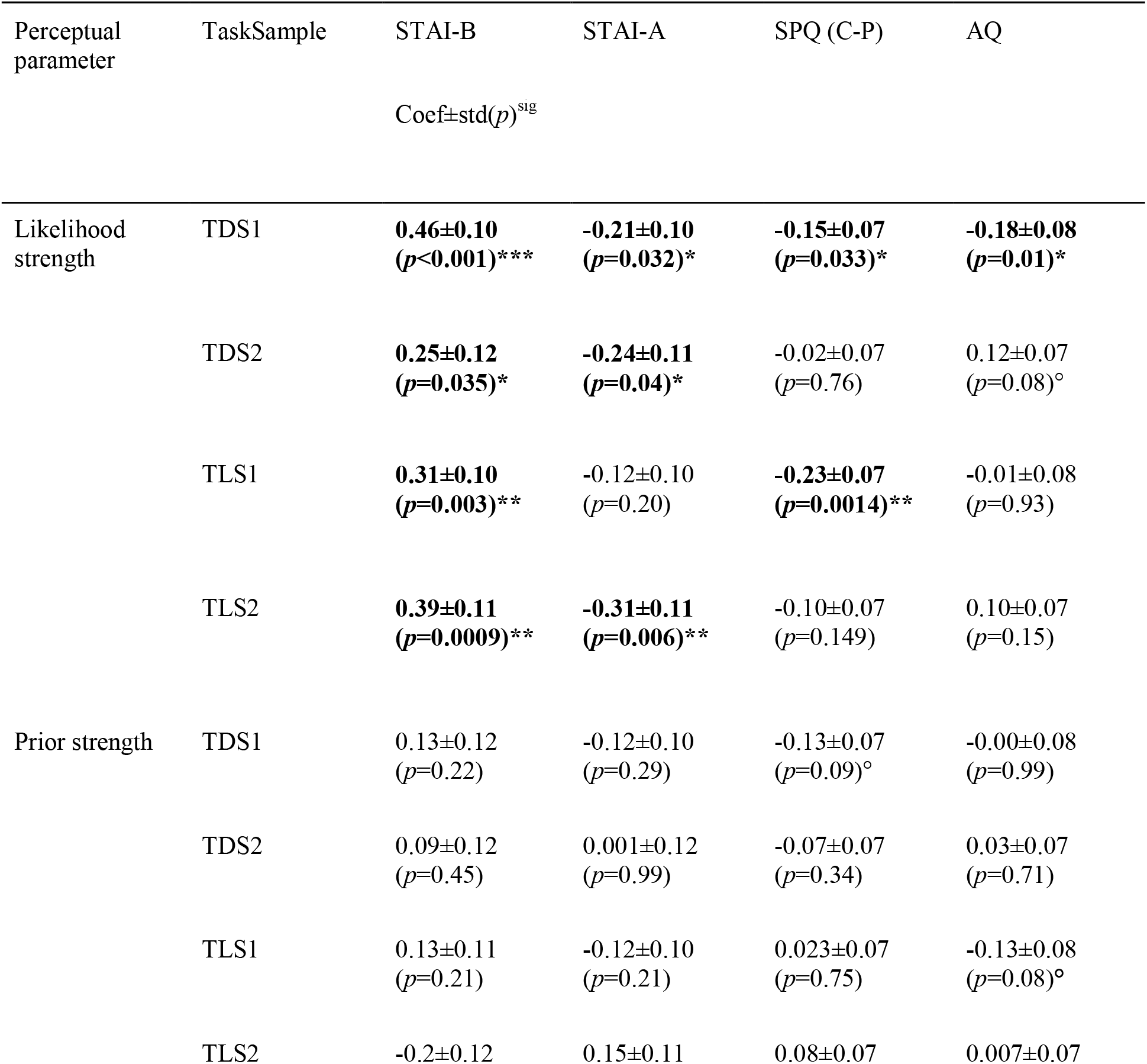
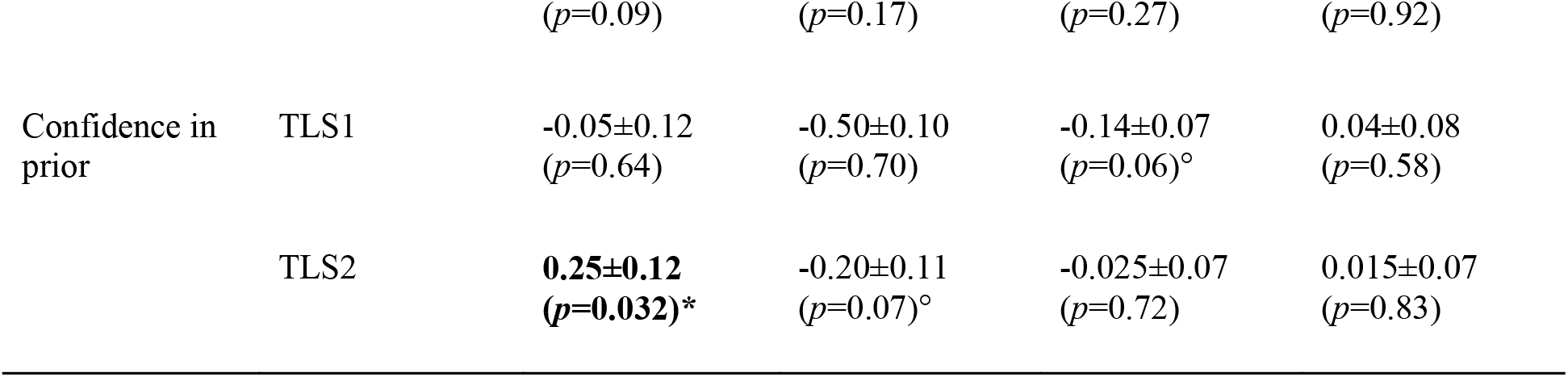
Coefficients of linear regression of response speed coefficients onto psychological traits. Each cell shows the coefficient ± SD and p-value for the trait predictor. For each subject, response speed coefficients were first estimated as the relationship between perceptual inference parameters (likelihood strength, prior strength, confidence in prior) and response speed. These per-subject speed coefficients were then regressed onto psychological traits using ordinary least squares (eq. 8), estimated separately for each perceptual parameter, task (TDS1, TDS2, TLS1, TLS2), and sample. Each model included STAI-A, STAI-B, SPQ (Cognitive-Perceptual dimension), and AQ as predictors, along with socio-demographic covariates (age, sex, education level, handedness) and their interactions with traits. ***p<0.001, **p<0.01, *p<0.05, °p<0.10.

| Perceptual parameter | TaskSample | STAI-B | STAI-A | SPQ (C-P) | AQ |
|---|---|---|---|---|---|
|  |  | Coef±std(*p*)<sup>sig</sup> |  |  |  |
| Likelihood strength | TDS1 | **0.46±0.10<br>(*p*<0.001)\*\*\*** | **-0.21±0.10<br>(*p*=0.032)\*** | **-0.15±0.07<br>(*p*=0.033)\*** | **-0.18±0.08<br>(*p*=0.01)\*** |
|  | TDS2 | **0.25±0.12<br>(*p*=0.035)\*** | **-0.24±0.11<br>(*p*=0.04)\*** | -0.02±0.07<br>(*p*=0.76) | 0.12±0.07<br>(*p*=0.08)° |
|  | TLS1 | **0.31±0.10<br>(*p*=0.003)\*\*** | -0.12±0.10<br>(*p*=0.20) | **-0.23±0.07<br>(*p*=0.0014)\*\*** | -0.01±0.08<br>(*p*=0.93) |
|  | TLS2 | **0.39±0.11<br>(*p*=0.0009)\*\*** | **-0.31±0.11<br>(*p*=0.006)\*\*** | -0.10±0.07<br>(*p*=0.149) | 0.10±0.07<br>(*p*=0.15) |
| Prior strength | TDS1 | 0.13±0.12<br>(*p*=0.22) | -0.12±0.10<br>(*p*=0.29) | -0.13±0.07<br>(*p*=0.09)° | -0.00±0.08<br>(*p*=0.99) |
|  | TDS2 | 0.09±0.12<br>(*p*=0.45) | 0.001±0.12<br>(*p*=0.99) | -0.07±0.07<br>(*p*=0.34) | 0.03±0.07<br>(*p*=0.71) |
|  | TLS1 | 0.13±0.11<br>(*p*=0.21) | -0.12±0.10<br>(*p*=0.21) | 0.023±0.07<br>(*p*=0.75) | -0.13±0.08<br>(*p*=0.08)° |
|  | TLS2 | -0.2±0.12<br>(*p*=0.09) | 0.15±0.11<br>(*p*=0.17) | 0.08±0.07<br>(*p*=0.27) | 0.007±0.07<br>(*p*=0.92) |
| Confidence in prior | TLS1 | -0.05±0.12<br>(*p*=0.64) | -0.50±0.10<br>(*p*=0.70) | -0.14±0.07<br>(*p*=0.06)° | 0.04±0.08<br>(*p*=0.58) |
|  | TLS2 | **0.25±0.12<br>(*p*=0.032)\*** | -0.20±0.11<br>(*p*=0.07)° | -0.025±0.07<br>(*p*=0.72) | 0.015±0.07<br>(*p*=0.83) |

In sum, the analyses of reaction times and choices lead to similar conclusions about the associations between psychological traits and interindividual differences in perceptual decision making. Stronger anxious traits are specifically associated with an increased effect of sensory likelihood, compared to the effect of the prior at the decision stage, or learning-related parameters; in contrast, associations with autistic and schizotypal traits were weaker or inconsistent across the two samples.

Associations between psychological traits and subject-specific speed coefficients derived from a multiple linear regression of speed on likelihood strength, prior strength, and confidence in prior simultaneously (one coefficient per predictor per subject). Each panel shows results from a separate multiple linear regression of one speed coefficient onto traits, demographics, and their interactions. Same plotting convention as in Fig. 4.

## 4. DISCUSSION

In this study, we characterised inter-individual differences in perceptual inference in the general population under the Bayesian framework with a multidimensional approach. We found that anxious, autistic and schizotypal traits correspond to three distinct, yet correlated dimensions. To overcome the limitations of previous studies, we used a set of two tasks with explicit and implicit priors. We combined these tasks with a model devised to differentiate the influence of priors from likelihood, and learning from decision-making, allowing us to quantify participant-specific parameters of perceptual inference. We found that subjects with more anxious traits performed better and relied more on sensory likelihood in their decision, with no difference in learning. No difference was consistently associated with autistic and schizotypal traits in the original and replication samples. These conclusions, derived from the analysis of choices, were corroborated by the analysis of response speed. In contrast to trait anxiety, state anxiety was associated with a reduced effect of sensory likelihood on response speed, an effect which was not captured in the analysis of choices.

These results are compatible with previous findings, as anxious traits are known to be associated with better academic achievement [82,83] and better performance in perceptual tasks [84,85]. Other Bayesian accounts have instead reported increased reliance on priors during perceptual decision-making associated with trait anxiety [46], which was not observed here. Interestingly, these results were obtained with priors which were fixed and stable across trials [46]. By contrast, in our task, priors varied over time and subjects, in particular anxious ones, may use perceptual priors differently in stationary and volatile environments [86]. State anxiety, unlike trait anxiety, showed a trend towards worse performance and a decreased effect of sensory likelihood on response speed. In line with this, previous studies have found that state anxiety often impairs performance and learning [87–89], especially in volatile environments [88,90]. Piray and Daw (2021) proposed that anxiety may result from a misinterpretation of stochasticity (random environmental noise) as volatility (meaningful changes in environmental structure) [44]. However, we did not find evidence for altered volatility estimates in our task.

Our findings that anxious traits in healthy individuals are associated with greater weight on sensory likelihood may reflect a milder, adaptive expression of mechanisms implicated in anxiety disorders. A moderate bias toward sensory evidence may sharpen perceptual decisions, while an excessive weighting could become maladaptive. Consistent with this view, several models propose that anxiety symptoms are driven by hypervigilance: an exaggerated allocation of attention to external sensory information, particularly threat-relevant information, at the expense of prior-based information that would otherwise attenuate anxiety (e.g., safety cues) [91]. Relatedly, interoceptive predictive-coding accounts propose that anxiety-prone individuals amplify bodily sensory signals and interpret them as threatening, so that small fluctuations generate disproportionate prediction errors, intensifying feared sensations and contributing to the somatic symptoms of anxiety [92–94].

Turning to autistic traits, we did not find any differences in perceptual inference. In a systematic review of 83 publications about the Bayesian hypothesis of autism [33], 2⁄3 of the studies did not find any differences in perceptual inference. The remaining 1⁄3 of the studies found a reduced influence of priors relative to sensory likelihood. A recent meta-analysis found a small-to-moderate overall effect in the same direction, but with significant unexplained heterogeneity across the 23 studies, thus offering only limited support for the Bayesian model of autism [35]. We did not reproduce this finding in our multidimensional analysis, but we found it in the unidimensional analysis of the replication sample, so that co-occurring anxiety - rather than autistic traits per se - might have driven previous results.

Concerning schizotypal traits, in the first sample, higher scores were associated with lower weight of sensory likelihood at the decision stage. This is consistent with evidence of stronger relative weighting of priors over sensory likelihood, thought to play a role in hallucinations [13,14,21– 27,95,96]. Conversely, our findings conflict with evidence of overweighted sensory evidence relative to priors, leading to perceptual instability and disordered salience attribution [12,29–32,97,98]. However, our result was not replicated in the second sample, leaving our conclusions tentative. We hypothesize that this lack of replication may reflect the significantly higher schizotypal cognitive-perceptual trait scores in the first sample, consistent with the view that alterations in perceptual inference differ qualitatively across levels of symptom severity [99–101].

Despite a methodological design developed to refine previous results, our study has several limitations. First, we used houses and faces as stimulus categories. Although these categories have been widely used in autism research, and some studies evidenced trait-related differences [102,103], altered face processing is common in autistic subjects [104] and might have masked effects of autistic traits on perceptual inference parameters. However, people with higher autistic traits did not perform worse, nor did they show face-related biases in learning and decision making, suggesting no particular alteration for this category of stimuli. Second, the timing of data collection may have affected sample characteristics. Indeed, the first sample was acquired shortly after the COVID-19 lockdown, when more people were at home and available for online studies, potentially recruiting participants less accustomed to cognitive tasks. Additionally, the psychological burden of the pandemic may have affected participants’ mental state and task engagement. Consistent with this, the proportion of subjects with aberrant responses was larger in the original sample (20%) than in the replication sample (5.4%). Third, our task assessed only stimulus categorisation. Our conclusions may not generalise to the estimation of continuous variables for which prior and likelihood weights cannot be distinguished [34].

Together, these findings suggest that anxious traits significantly modulate perceptual inference with better performances in categorising noisy stimuli thanks to a greater reliance on sensory likelihood at the decision stage. They underscore the value of multidimensional approaches to take into account the intercorrelation of psychological traits and the advantage of using tasks able to distinguish learning and decision processes involved in perceptual inference. Future studies using our design in psychiatric populations may further elucidate how perceptual inference is altered in patients as a function of diagnosis and clinical transdiagnostic dimensions such as anxiety, autistic and psychotic traits, refining the relationship between symptom expression and cognitive impairment.

## 5. SUPPLEMENTARY MATERIALS

### 5.1. Supplementary Methods

#### 5.1.1. Pre-registration

A preregistration was conducted for the study before the acquisition of the second sample (https://osf.io/k9vqd).

#### 5.1.2. Tasks

The tasks were run on GorillaTM. The experiment was composed of two tasks: the decision task (with explicit priors) and the decision-and-learning task (with implicit priors), in this order. They contained respectively 300 trials and around 600 trials (mean=597, SD=15; variability arises from sampling the length of each stable period, see below). Each task was divided into three parts separated by self-paced breaks. Stimuli were noisy, gray-scaled, morphed images of faces and houses. Noise was first added on images with the GNU image. Morphed images were created in the Fourier space, by computing a weighted average of the phase values of each pair of face and house images, and using the mean amplitudes of all images (all images therefore have the same frequency spectrum). The level of perceptual evidence in favour of the house/face category was estimated empirically for each image as the proportion of house/face responses in a categorisation task performed by a group of 47 typical human observers who did not participate in the main experiment, see [78]. Each trial of the main experiment consisted in the categorization of an image presented for 150ms. Subjects reported their responses on a keyboard with the keys ‘e’ and ‘p’ for face and house (counterbalanced across subjects), within a response window of 2000ms. In the decision task, each image was preceded by a cue consisting of a set of pictograms. Indicating the prior probability of the house/face category, presented for 1500 ms. In the decision-and-learning task, the latent prior value was stable in periods of 40 trials (±4) separated by unsignaled change point. In both tasks and on each trial, the latent category was sampled according to its prior probability, and an image corresponding to this category was sampled pseudo-randomly. To make the detection of a change point easier in the decision-and- learning task, three images with strong likelihood (p(house)>0.85 or <0.15) were presented among the first four images after each change point. In the decision task, subjects were prompted to report the prior value corresponding to each pictogram at the end of the task on a quasi-continuous rating scale. In the decision-and-learning task, subjects were periodically (every 14 trials) asked to report the current (implicit) prior value. We used a cover story to gamify the task and make it intuitive (Figure S1, S2).

**Figure S1.**
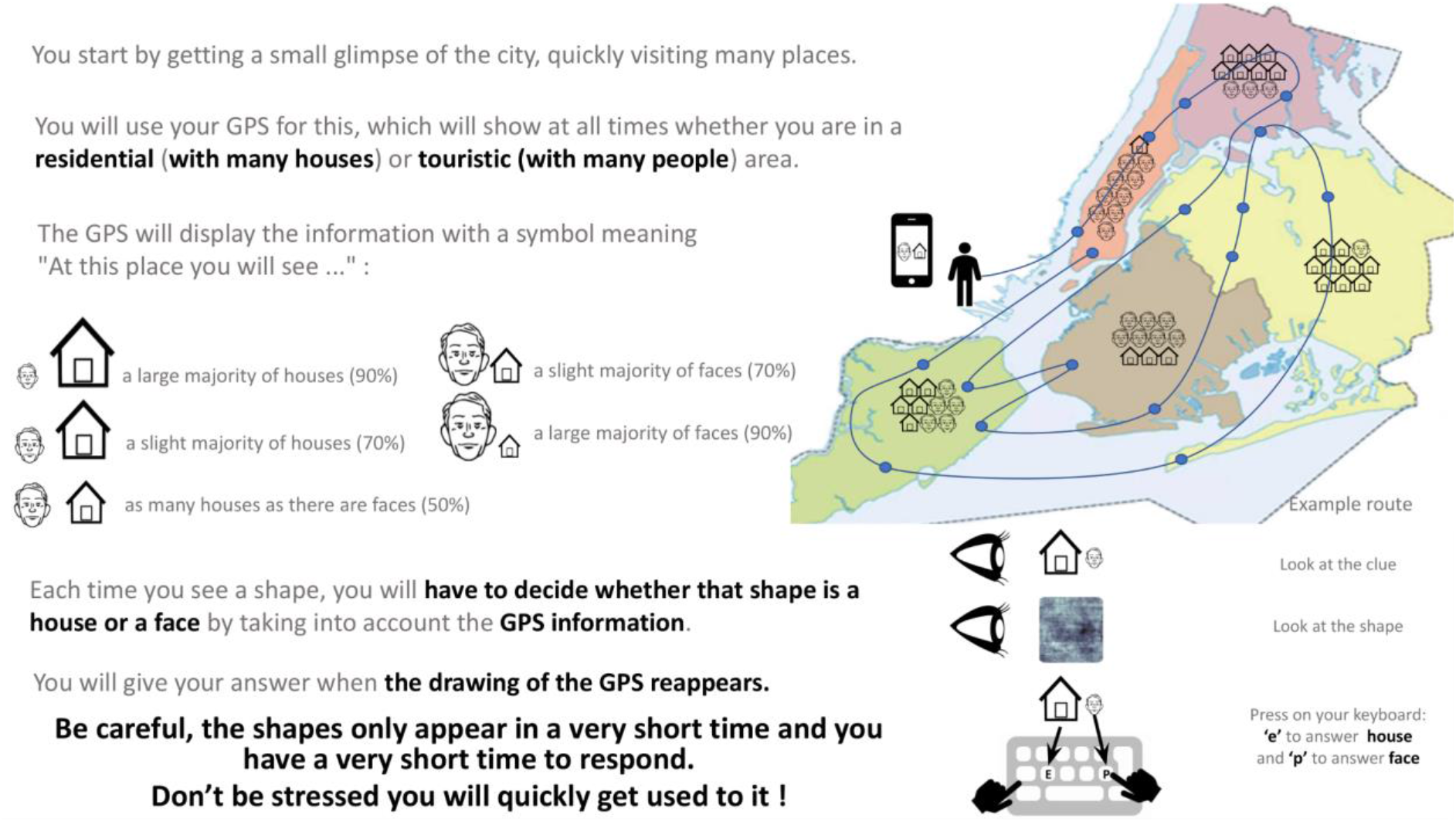
Instruction slide for the decision-only task with explicit priors.

**Figure S2.**
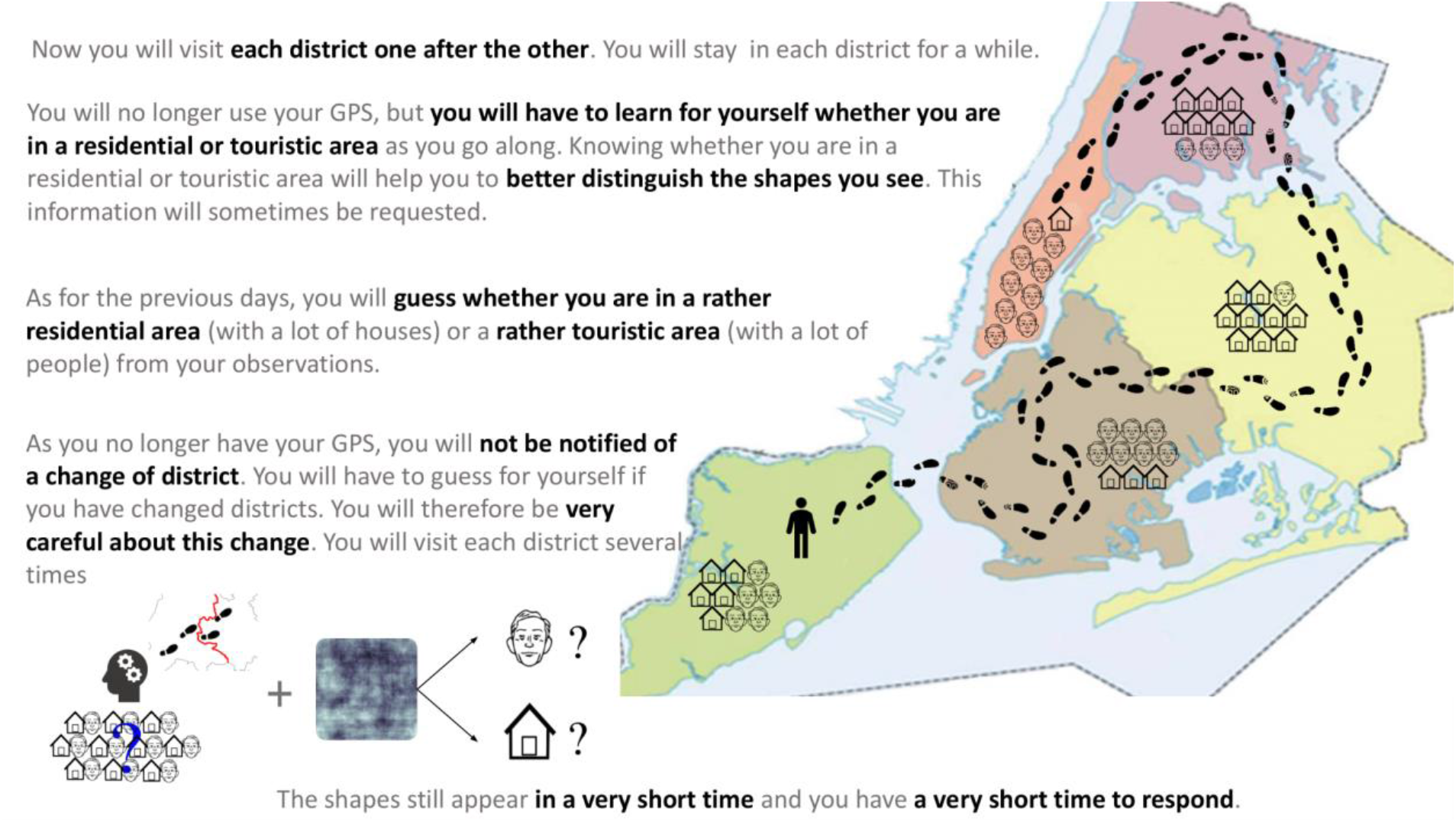
Instruction slide for the decision-and-learning task with implicitly learnt priors.

#### 5.1.3. Computational models

The models used in this study were introduced in Bévalot & Meyniel, 2024 [78]. Instead of referring the reader to this publication, we reproduced here the description of the models.

##### Bayesian perception model

Both tasks are to infer, on each trial k, whether the latent category c_k_ is a house (denoted *H*) or a face (*F*) based on the noisy image I_k_ and some prior about the category. We focus first on the decision task, in which the prior value does not depend on previous images, but only on the cue. Formally, this inference amounts to estimating the posterior probability *p*(*c_k_*=*H*|*I_K_*), which is done optimally with Bayes rule:

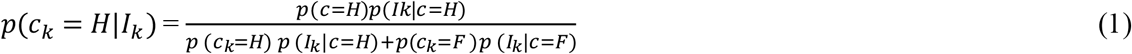

where *p*(*c_k_*=*H*) is the prior probability of the latent category being a house on trial *k*, which we note *θ_k_* for brevity below (note that *p*(*c_k_*=*F*) =*1* −*θ_k_* and *p*(*c_k_*=*H*|*I_K_*) is the sensory likelihood about the house category based on image *I_k_* only. We introduce the notation 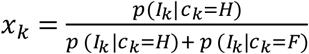 for the normalized sensory likelihood, and rewrite equation (1) more concisely as:

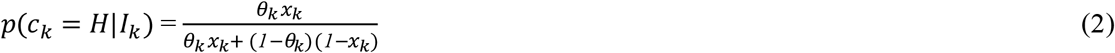

The equation is simpler when using the log-odd transformation 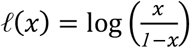:

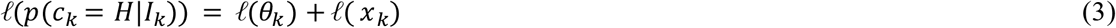

The (normalized) sensory evidence *x_k_* corresponding to each image was estimated empirically in an independent group of subjects.

The prior *θ_k_* was equal to its explicit value in the decision task. To analyse the decision-and-learning task, we used in different analyses either its generative value, or its value learned based on previous images by the BAYES-OPTIMAL model or the BAYES-FIT-ALL model.

##### BAYES-OPTIMAL learning model

The BAYES-OPTIMAL observer learns the prior θ by updating optimally its posterior distribution after every image in the sequence, starting with a uniform distribution before the first image. The generative process of the sequence has an interesting property known as the Markov property: if the previous prior value *θ_k-1_* is given, then the previous observations *I_1_*,…, *I;_k-1_* (denoted *I_1: k−1_* below) are not informative about the current prior value *θ_k_*. This is because *θ_k_* will be equal to *θ_k−*1*_* if no change point occurs; change points occur with probability *v* (called volatility) otherwise, and this does not depend on *I_1: k−1_*:. This Markov property makes it possible to cast the updating process as the forward pass of a hidden markov chain [105–107], resulting in the following iterative equation for learning the prior θ:

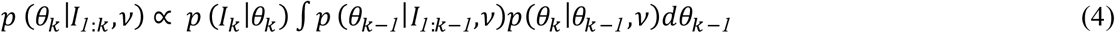

where *p (I_k_ |θ_k_)* is the likelihood of the current image given some prior value *θ_k_*; which is obtained by marginalizing over the unknown latent category:

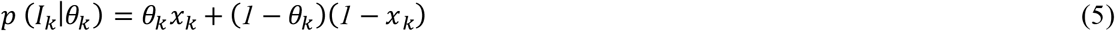

We computed the integral in eq. 4 numerically by discretization on a grid. Note that eq.4 returns the inferred prior value after having observed *I_k_*. In eq. 2, the prior that is used to infer the category of image *I_k_* is the prior estimated before seeing *I_k_*; it is obtained by marginalizing over the values of *θ_k-1_* given the previous observations *I_1 k;-1_*:

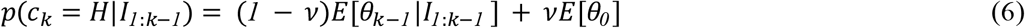

where E[·] denotes the expectation and *θ_0_* is the distribution from which new values of θ are sampled in case a change point occurs.

##### BAYES-FIT-ALL model

This model differs from the BAYES-OPTIMAL with 6 free parameters that are fit to the choices of each participant. At the learning stage (eq.4), the volatility parameter is a free parameter because the value assumed by participants may differ from the generative value. This BAYES-FIT-ALL model also allows for distortions of the likelihood function *p*(*I_k_* |*θ_k_* that can exacerbate it, dampen it, or bias it. We considered a distortion that is affine in log-odd [108], hence yielding two additional free parameters. At the decision stage, the BAYES-FIT-ALL model has three free parameters because it considers subject-specific weights on the likelihood *w_DL_* and prior *w_DP_* terms in eq.3 and an additive bias term *δ_D_* (in the BAYES-OPTIMAL model, *w_DL_* = *w_DP_*= 1 and *δ_D_* = 0). The resulting modified eq.3 can be fit to each participant’s choices with a logistic regression:

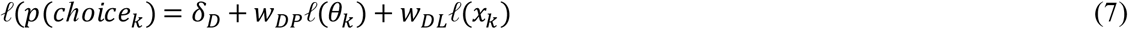

We fitted the model parameters for each subject separately, by minimizing the negative log-likelihood of the subject’s choices given the parameterized model with L_2_ (ridge) penalty. The ridge penalty was not applied on the parameters themselves since several of them are normatively expected to have non zero values (*v,w_DL_,w_DP_,w_LL_*), instead it was computed on the difference between fitted and optimal parameters, with a penalty set to 0.1 (this value was selected among 0.001, 0.01, 0.1, 1, 10 because it prevented occasional aberrant parameter values while maintaining high log-likelihood values). Minimization was performed with the Powell method implemented in Scipy (Python), retaining the best fit after 100 random initializations of the minimizer. Parameter recovery was satisfactory (Figure S1).

#### 5.1.4. Statistical methods

Wilcoxon tests were used when data were not normally distributed. Bootstrapped differences between regression coefficients were used to test the specificity of trait associations across dependent variables (see below).

##### Traits analysis

For each psychometric scale (STAI, SPQ, AQ), we computed Crohnbach’s alpha, evaluating the internal consistency of answers provided to the different items. Cronbach’s alpha analysis evaluates the internal consistency of a psychometric scale by calculating how well the items on the scale correlate with each other, reflecting whether they measure the same underlying construct. It produces a coefficient between 0 and 1, where higher values indicate stronger internal consistency and better reliability of the scale. We used the Python package pingouin-stats [109]. We analysed the underlying dimensions covered by the items of the different scales with a factor analysis using the FactorAnalyze Python package. (https://github.com/EducationalTestingService/factor_analyzer). We set the number of factors to 4 to examine whether the three dimensions from our questionnaires corresponded to distinct, identifiable factors, or if introducing an additional factor would alter the grouping of items.

##### Dominance analysis

To measure the strength of the modulation of each explanatory variable on the response variables, we ran a dominance analysis. A dominance analysis involves regressions using every possible combination of explanatory variables in sub-sets. The total dominance of a given explanatory variable is the average difference in the coefficient of determination (R^2^) when this explanatory variable is added to the set and subsets of the other explanatory variables. (see https://github.com/dominance-analysis/dominance-analysis for more details)

##### Multiple linear regression models to quantify associations with traits

We used as response variables *y_i_* the parameters of perceptual inference (e.g. the weights of priors in decision, *w_DP_*) obtained from each subject *i*. The explanatory variables were the psychological traits ***T*** (scores from STAI-A, STAI-B, AQ, SPQ Positive), and the socio-demographic data ***C*** (age, sex, education level, handedness). Each multiple linear regression included interactions between psychological traits and demographic data:

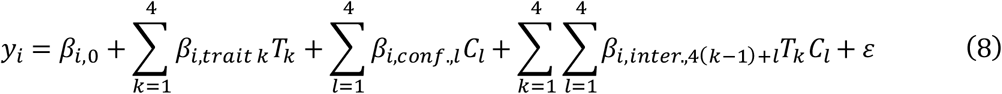

Separate linear models were estimated for each parameter of perceptual inference, each task (decision or learning-and-decision) and each sample (original, replication). These linear models were estimated using ordinary least squares (OLS). We ran a similar analysis with each of the fitted parameters as dependent variables.

We also analysed the effects of the likelihood strength, the prior strength, and confidence about the prior value onto response speed (the inverse of response time) in linear regressions (one regression per factor), and we used the linear coefficients as response variables in (8). We quantified the likelihood strength of the image presented at trial k as:

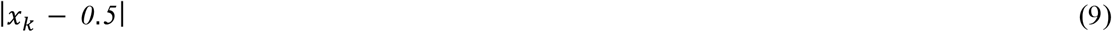

The prior strength, reflecting how strongly one can predict the next image category at trial *k* based on the value of the prior as:

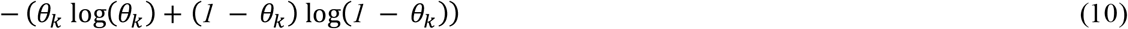

Note that formally, Eq. 10 corresponds to Shannon entropy, a principled measure of how informative the prior is about the next image category.

The confidence about the prior value on trial k as the log precision of its posterior distribution (Eq. 4):

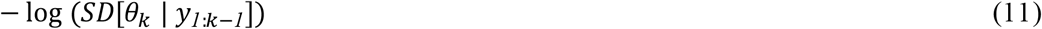

Regressions were fit using scikit-learn’s LinearRegression module [110] to run the linear regressions.

##### Bootstrap Regression Analysis

We compared trait effects across perceptual parameter weights using bootstrap resampling. For each trait, we fit two OLS models predicting two different weights, with z-scored predictors and traits x demographic interactions. The standardized trait coefficient was extracted from each model, and their difference (Δ) served as the point estimate. The bootstrap distribution of Δ was obtained via 1000 resamples with replacement, refitting both models on each resample. The 95% CI was defined as the 2.5th and 97.5th percentiles of this distribution. A CI excluding zero was taken as evidence of a meaningful difference in trait effects between the two weights.

#### 5.1.5. Data availability

The raw data will be made available on OSF upon publication.

#### 5.1.6. Code availability

The code to reproduce the results will be made available on OSF upon publication.

### 5.2. Supplementary Results

#### Supplementary Tables

**Table S1.** Internal consistently of questionnaires using Cronbach’s alpha. Results are presented as alpha[ci].

|  | S1 | S2 |
|---|---|---|
| STAI-A | 0.93[0.92-0.94] | 0.94[0.93-0.95] |
| STAI-B | 0.93[0.92-0.94] | 0.95[0.94-0.95] |
| AQ | 0.79[0.76-0.83] | 0.83[0.80-0.85] |
| SPQ Total | 0.94[0.930.95] | 0.94[0.93-0.95] |
| SPQ Positive | 0.90[0.89-0.92] | 0.89[0.87-0.91] |
| SPQ Negative | 0.88[0.86-0.90] | 0.91[0.89-092] |
| SPQ Disorganization | 0.87[0.85-0.89] | 0.86[0.83-0.88] |

**Table S2.** Coefficients of linear regression of perceptual inference parameters (logistic regression weights) onto psychological traits. Each cell shows the coefficient ± SD and p-value for the trait predictor. Models were estimated separately for each perceptual parameter (weights of posterior, likelihood, prior), task (TDS1, TDS2, TLS1, TLS2), and sample, using ordinary least squares (eq. 8). Each model included STAI-A, STAI-B, SPQ cognitive-perceptual dimension, and AQ as predictors, along with socio-demographic covariates (age, sex, education level, handedness) and their interactions with traits. ***p<0.001, **p<0.01, *p<0.05, °p<0.10.

| Perceptual parameter | TaskSample | STAI-B<br>Coef±std(*p*)<sup>sig</sup> | STAI-A | SPQ (C-P) | AQ |
|---|---|---|---|---|---|
| Posterior (A) | TDS1 | 0.19±0.10 (*p*=0.064)° | -0.16±0.09 (*p*=0.085)° | **-0.25±0.07 (*p*=0.0002)\*\*\*** | -0.03±0.07 (*p*=0.73) |
|  | TDS2 | 0.15±0.12 (*p*=0.21) | -0.05±0.11 (*p*=0.65) | 0.02±0.07 (*p*=0.77) | 0.11±0.07 (*p*=0.12) |
|  | TLS1 | **0.26±0.10 (*p*=0.010)\*\*** | -0.16±0.09 (*p*=0.088)° | **-0.30±0.07 (*p*<0.001)\*\*\*** | 0.01±0.07 (*p*=0.91) |
|  | TLS2 | **0.25±0.12, (*p*=0.039)\*** | -0.09±0.12 (*p*=0.44) | 0.06±0.07 (*p*=0.37) | 0.02±0.07 (*p*=0.82) |
| Likelihood (B) | TDS1 | **0.30±0.11, (*p*=0.0054)\*\*** | -0.17±0.10 (*p*=0.086)° | **-0.22±0.07 (*p*=0.0029)\*\*** | -0.03±0.08 (*p*=0.73) |
|  | TDS2 | **0.24±0.12, (*p*=0.037)\*** | -0.12±0.11 (*p*=0.30) | -0.07±0.07 (*p*=0.35) | 0.13±0.07 (*p*=0.062)° |
|  | TLS1 | **0.25±0.10, (*p*=0.013)\*** | -0.14±0.09 (*p*=0.13) | **-0.33±0.07 (*p*<0.001)\*\*\*** | 0.04±0.07 (*p*=0.61) |
|  | TLS2 | **0.28±0.12 (*p*=0.018)\*** | -0.13±0.11 (*p*=0.25) | 0.05±0.07 (*p*=0.47) | 0.05±0.07 (*p*=0.50) |
| Prior (C) | TDS1 | -0.08±0.10 (*p*=0.44) | 0.02±0.10 (*p*=0.87) | **-0.15±0.07 (*p*=0.036)\*** | -0.03±0.08 (*p*=0.67) |
|  | TDS2 | -0.18±0.11 (*p*=0.12) | 0.11±0.11 (*p*=0.31) | 0.08±0.07 (*p*=0.22) | -0.003±0.07 (*p*=0.97) |
|  | TLS1 | 0.18±0.10 (*p*=0.083)° | -0.13±0.10 (*p*=0.18) | -0.08±0.07 (*p*=0.26) | -0.04±0.08 (*p*=0.57) |
|  | TLS2 | 0.02±0.12 (*p*=0.86) | 0.06±0.11 (*p*=0.62) | 0.05±0.07 (*p*=0.50) | -0.03±0.07 (*p*=0.72) |

**Table S3.** Standardised regression coefficients relating perceptual parameters to response speed (1/RT), by task and sample. For each subject, a multiple linear regression predicted z-scored response speed from z-scored likelihood strength and prior strength (TDS1, TDS2, TLS1, TLS2), with confidence in the prior (posterior log-precision inferred by the Bayes-optimal model) included as an additional predictor for the decision-and-learning tasks (TLS1, TLS2). Trials with RT < 100 ms or > 1950 ms were excluded prior to fitting. Regression coefficient columns report the mean±SD of per-subject coefficients across the sample; t(p) columns report one-sample t-tests of these coefficients against zero.

| Perceptual parameter | TaskSample | Regression coefficient (mean±SD) | T-test against 0 t(p) |
|---|---|---|---|
| Likelihood strength | TDS1 | 0.12±0.14 | 14.47 (p<0.0001) |
|  | TDS2 | 0.17±0.13 | 21.16 (p<0.0001) |
|  | TLS1 | 0.14±0.13 | 17.39 (p<0.0001) |
|  | TLS2 | 0.20±0.14 | 24.25 (p<0.0001) |
| Prior strength | TDS1 | 0.08±0.09 | 13.35 (p<0.0001) |
|  | TDS2 | 0.118±0.09 | 20.11 (p<0.0001) |
|  | TLS1 | 0.06±0.14 | 6.93 (p<0.0001) |
|  | TLS2 | 0.08±0.13 | 10.40 (p<0.0001) |
| Confidence in prior | TLS1 | 0.07±0.10 | 11.16 (p<0.0001) |
|  | TLS2 | 0.08±0.10 | 13.45 (p<0.0001) |

## Supplementary Figures

**Figure S3.**
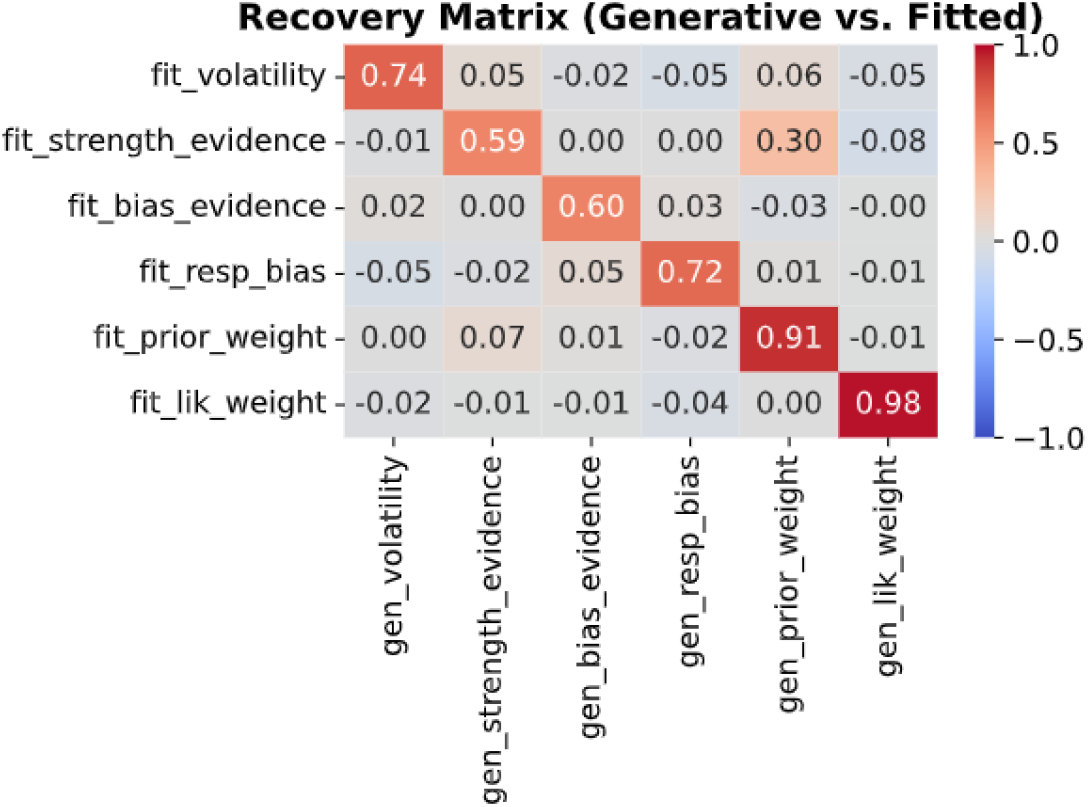
Parameter recovery for the BAYES-FIT-ALL model. For each subject, artificial behaviour was simulated using randomly drawn generative parameters, and the model was then fit to recover those parameters. The matrix shows Pearson correlations between generative and fitted values for each parameter. High correlations along the diagonal and near-zero off-diagonal correlations indicate that all parameters are identifiable and well recovered, validating that the fitting procedure returns meaningful parameter estimates. Decision parameters were recovered more accurately (likelihood weight r = 0.98, prior weight r = 0.91, response bias r = 0.72) than learning parameters (strength of evidence r = 0.59, bias of evidence r = 0.60, volatility r = 0.74).

**Figure S4.**
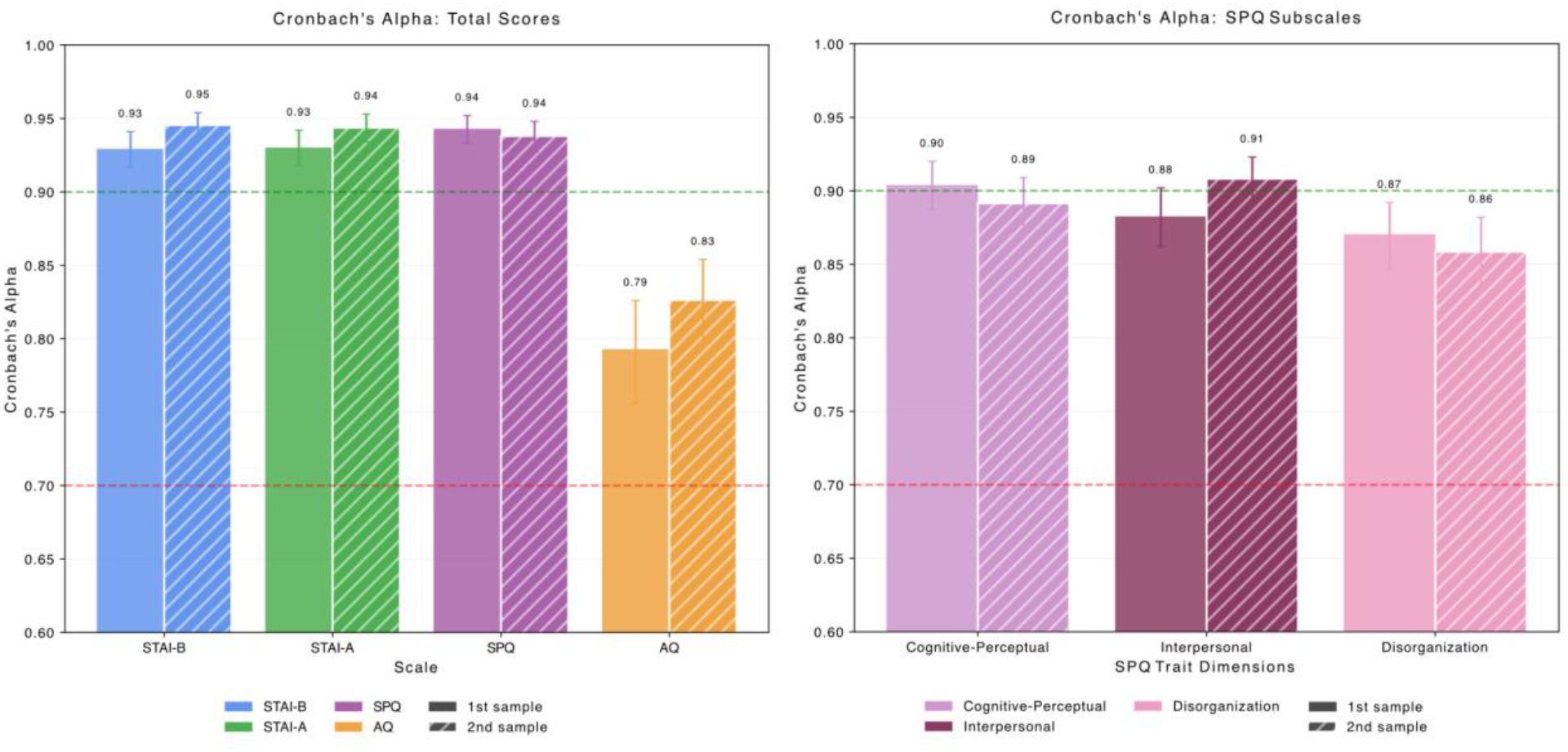
Cronbach’s alpha coefficients showing internal consistency of all questionnaires across samples S1 (Original) and S2 (Replication). Error bars represent 95% confidence intervals around each alpha coefficient. Solid bars indicate S1 data, while hatched bars indicate S2 data. Color coding represents questionnaire type: green for STAI-A, blue for STAI-B, orange for AQ, and pink for SPQ (with lighter pink and thinner bars for SPQ subscales). Horizontal dashed lines indicate acceptable (α = 0.70) and good (α = 0.90) reliability thresholds.

**Figure S5.**
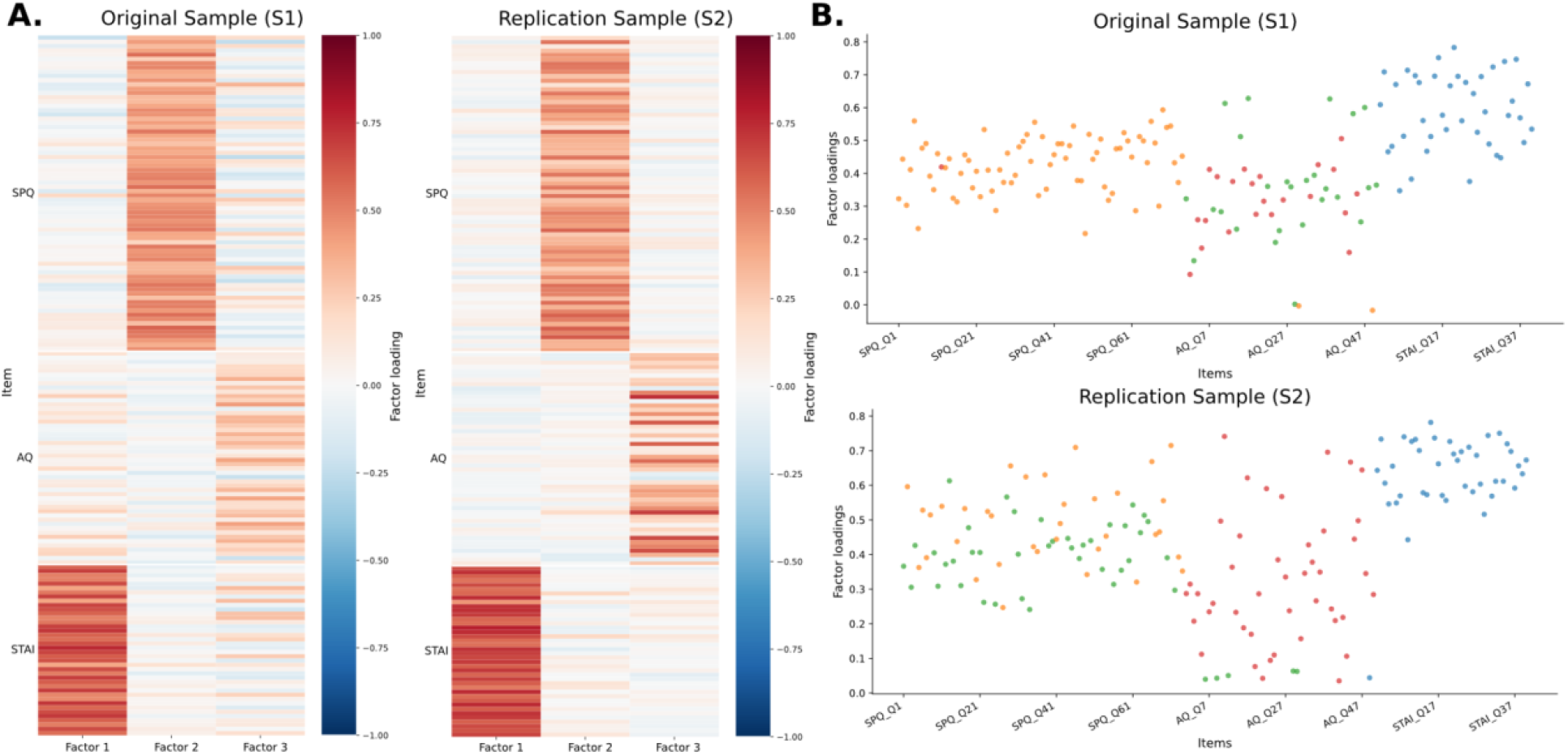
Factor loadings: anxious, autistic, and schizotypal traits form three distinct dimensions. Before adopting a multidimensional approach, we tested in a data-driven manner whether the three traits actually correspond to three different psychological dimensions using a factor analysis. Panel A: We found that a given factor typically corresponds to the items of a given questionnaire. Panel B: When four factors are used in the analysis (one more than the number of questionnaires), the fourth factor does not correspond to items from different scales (thus, a fourth dimension), instead, one questionnaire splits into two factors, that is, the AQ splits in S1, and the SPQ splits in S2. Therefore, the STAI, AQ and SPQ questionnaires correspond to three separable dimensions, and thus, traits.

**Figure S6.**
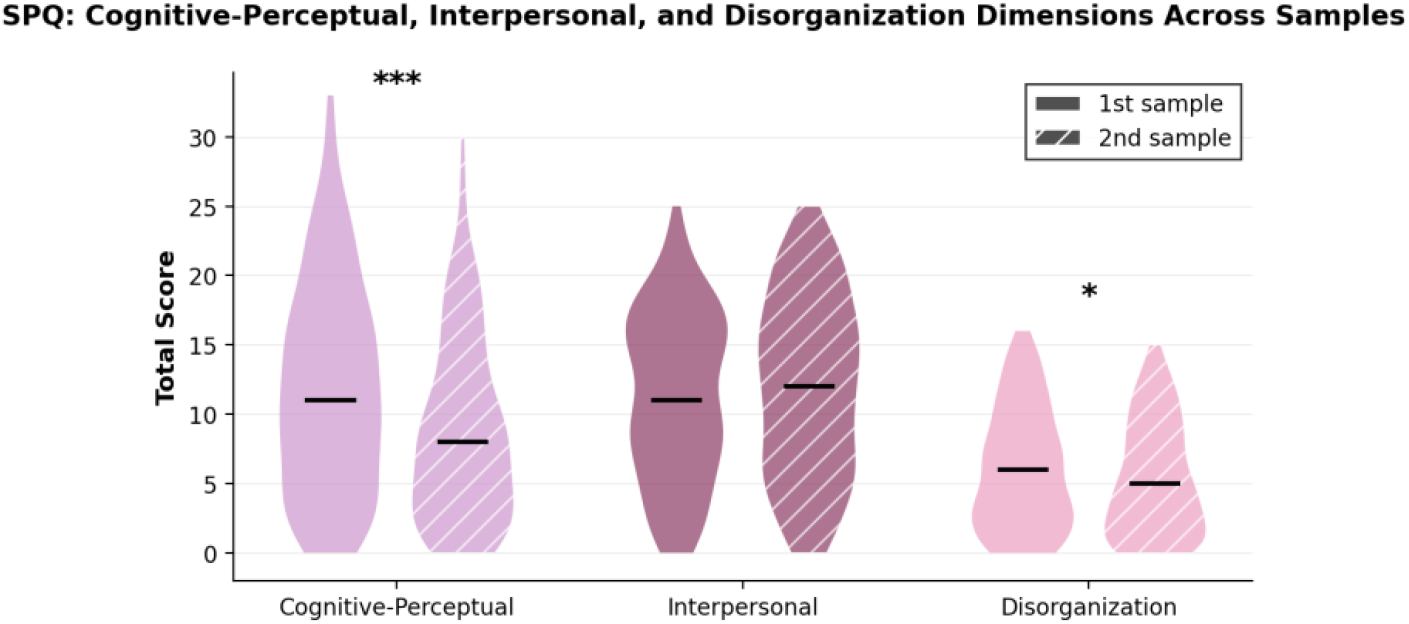
Distribution of SPQ subscales (cognitive-perceptual, interpersonal, and disorganisation) across samples S1 and S2. The original sample (S1) had significantly higher cognitive-perceptual traits (S1 mean = 11.89(7.63), S2 mean = 8.79(6.64); t-test: t = 5.11, p < 0.0001), and disorganization traits (S1 mean = 6.04(4.36), S2 mean = 5.27(4.13); t-test: t = 2.12, p = 0.03) compared to the replication sample (S2). Samples did not show any significant differences in interpersonal trait scores (S1 mean = 11.27(6.03), S2 mean = 11.95 (6.55); t-test: t = −1.27, p = 0.20).

**Figure S7.**
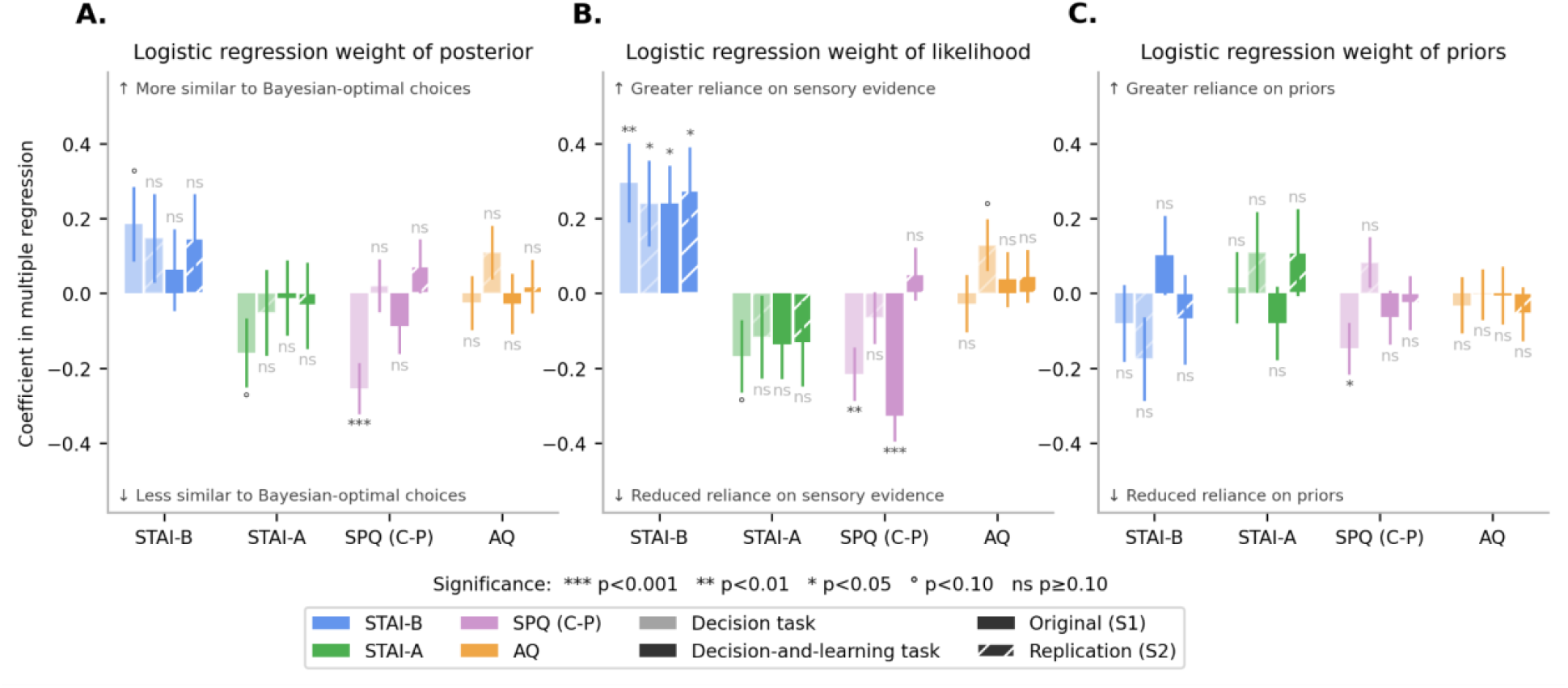
Results of linear regression of traits onto logistic regression weights of posterior, likelihood and priors in the decision and learning task with BAYES-FIT-ALL fitted priors.

**Figure S8.**
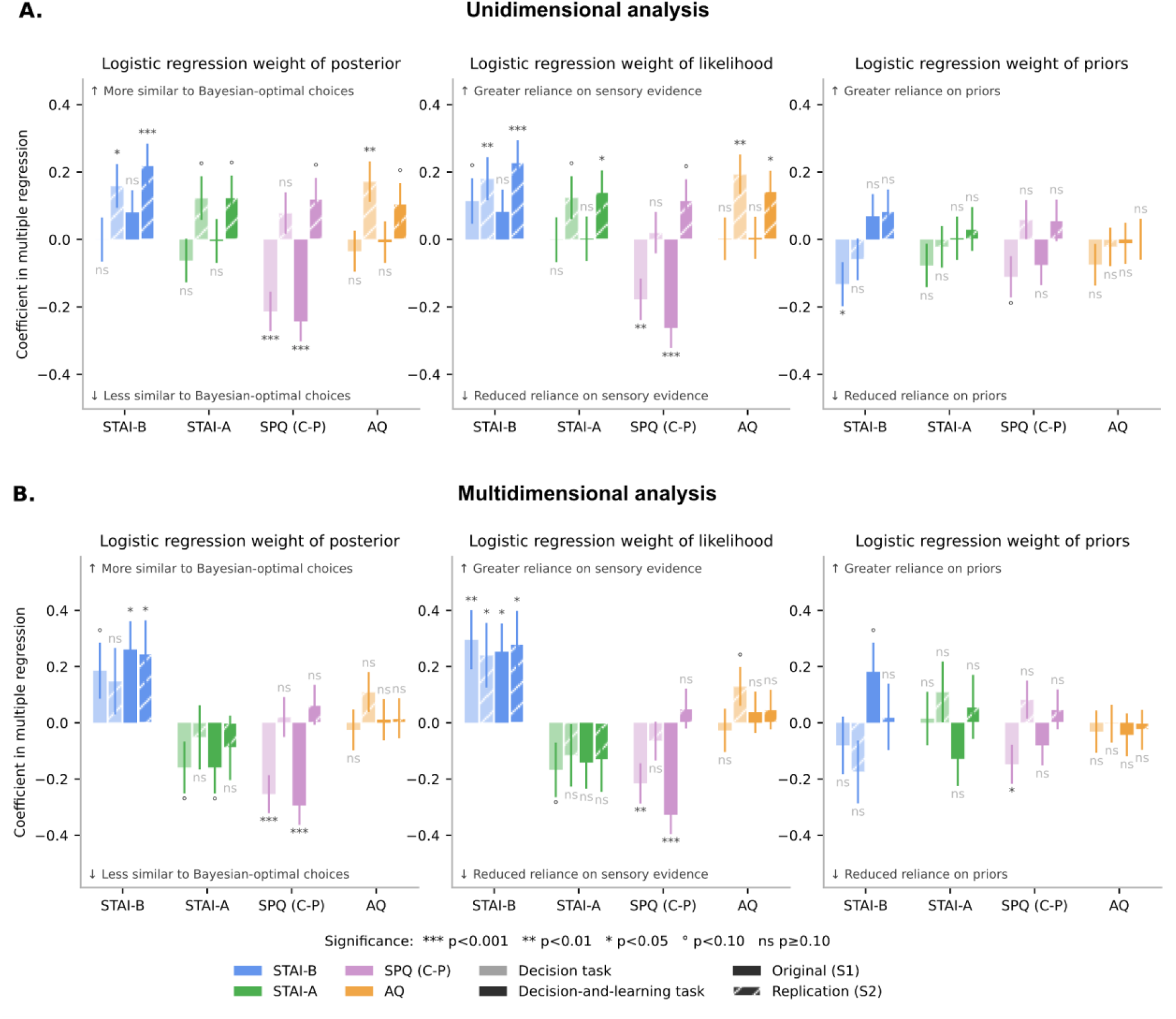
Linear regression of logistic regression weights onto traits using a (A) unidimensional approach versus a (B) multidimensional approach A unidimensional approach could miss trait-specific effects, or spuriously detect an effect that is actually due to another dimension. Here, we see that a unidimensional analysis would have concluded that autistic traits are associated with a difference in the weight of sensory likelihood (TDS1=−0.0043±0.063 (p=0.94), TLS1=−0.0061±0.063 (p=0.92), TDS2=−0.16±0.058 (p=0.0044), TLS2=0.10±0.051 (p=0.046)) which vanishes in a multidimensional analysis (below).

**Figure S9.**
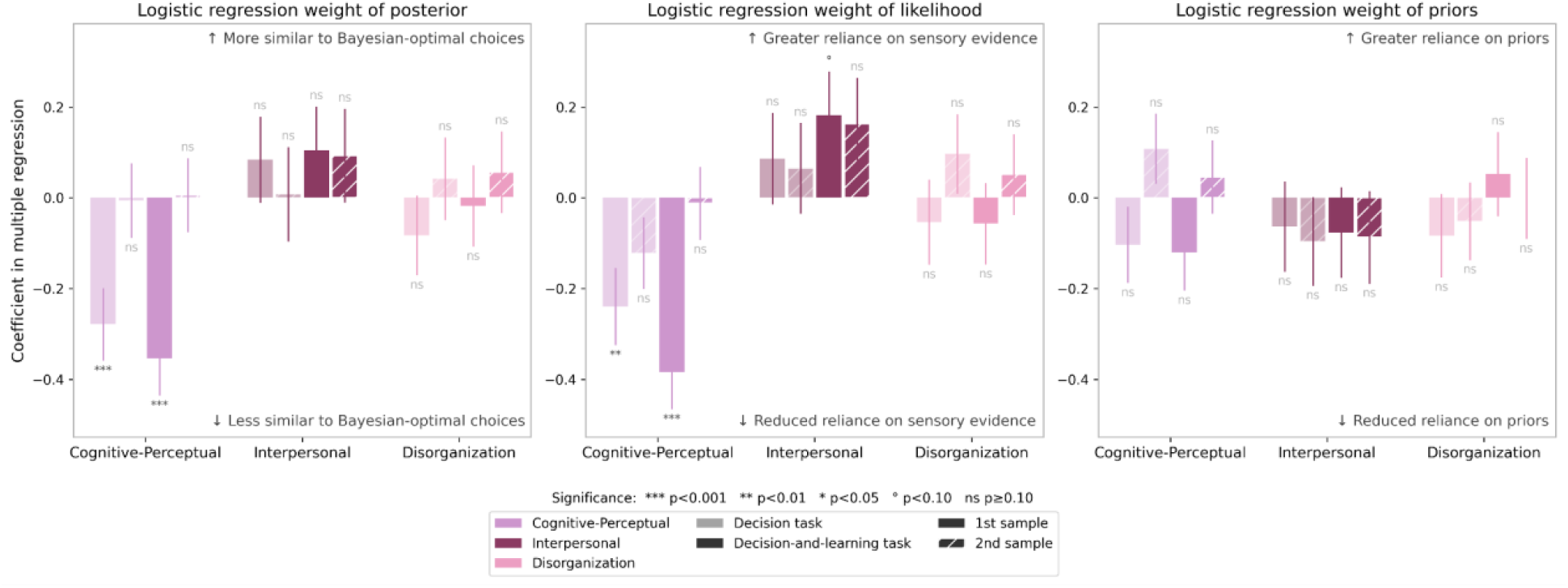
Linear regression of SPQ trait dimensions(cognitive-perceptual, interpersonal, disorganization) onto logistic regression weights of posterior, likelihood and priors in the decision and learning task, using optimal priors. Linear regressions also include other traits and demographic variables, but we only show SPQ dimensions. The cognitive-perceptual dimension is associated with reduced weight of sensory evidence and choices further from the Bayes optimal posterior. Interpersonal and disorganization symptoms show no significant impact on sensory processing.

**Figure S10.**
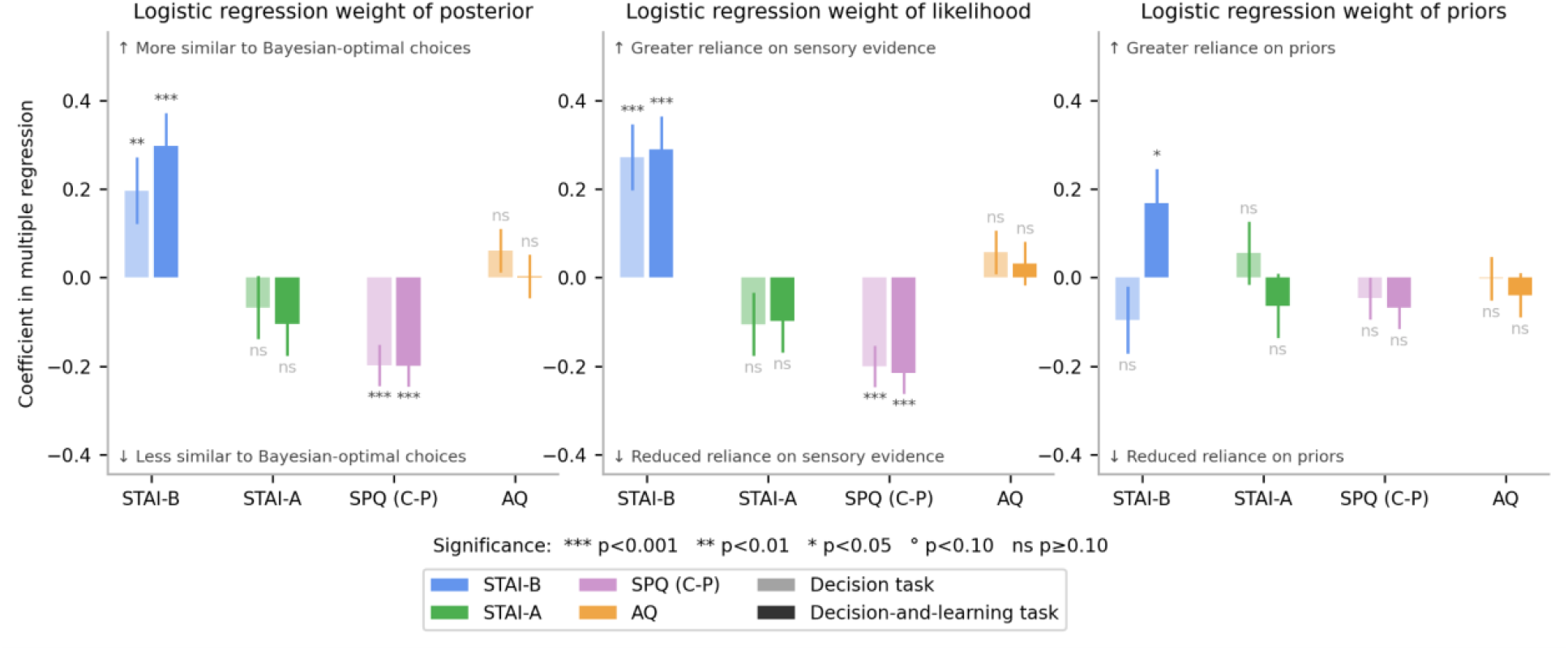
Linear regression of traits onto logistic regression weights of posterior, likelihood and priors in the decision and learning task, using optimal priors with samples S1 and S2 merged.

**Figure S11.**
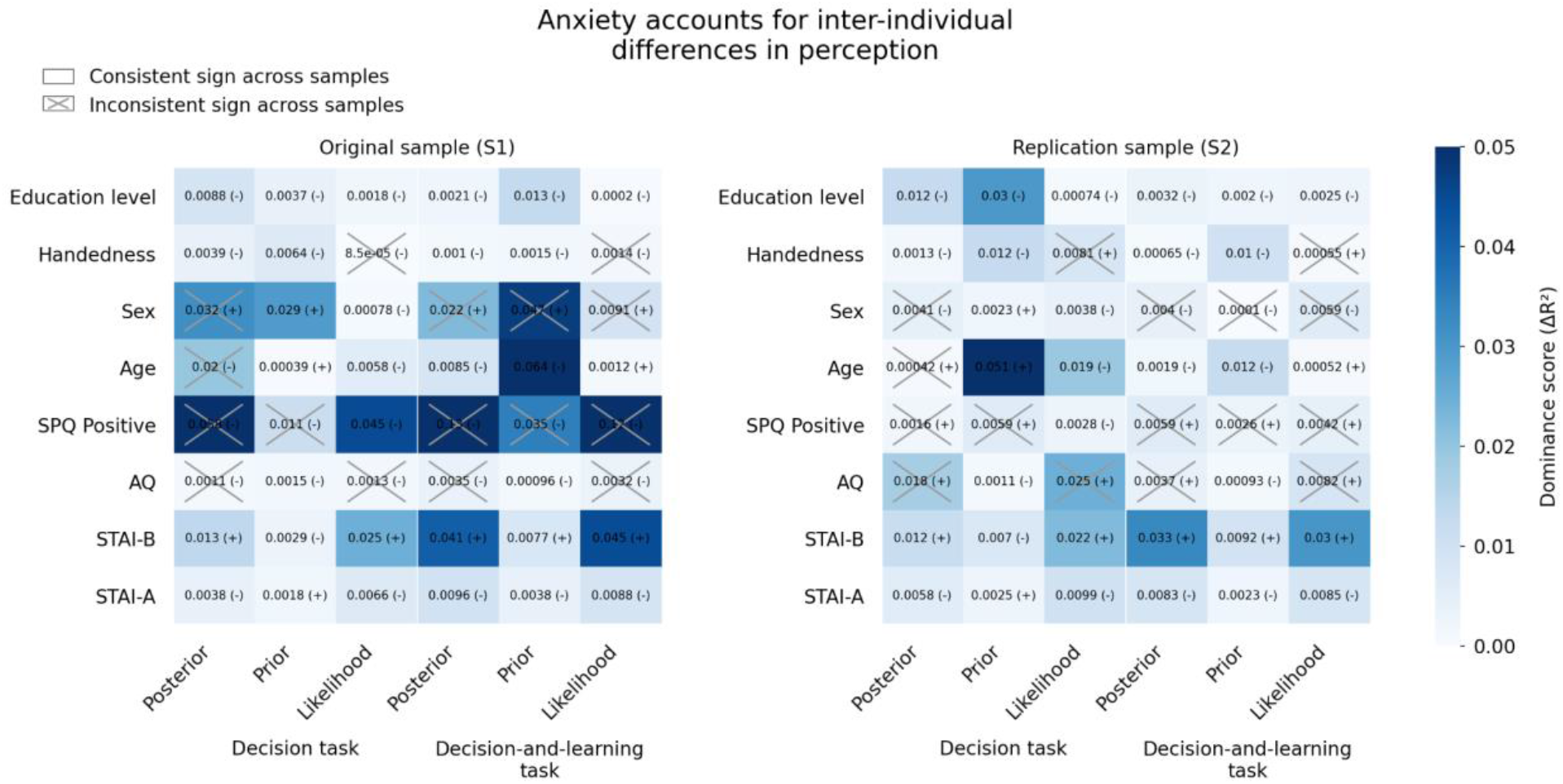
Dominance analysis identifies associations between perceptual inference and anxiety. Dominance scores were computed across different features (rows) separately for each parameter of perceptual inference (columns), and separately for the two samples. Total dominance corresponds to the mean incremental proportion of variance explained (ΔR^2^) by each feature across all possible subsets of features. Effects that were inconsistent between samples (i.e., opposite signs) are indicated with crosses. Effects correspond to regression coefficients obtained from models including all features simultaneously (no interactions), fitted separately for each perceptual parameter and sample.

## Notes

### Competing Interest Statement

The authors have declared no competing interest.

### Summary of Updates

We improved the fitting and the regression method to improve the robustness of our results

